# Investigating dietary microRNA stability and function using a transgenic milk model with unique microRNA sequences

**DOI:** 10.64898/2026.04.13.718217

**Authors:** Zeinab Husseini, Nathalie Majeau, Ismail Fliss, Abderrahim Benmoussa

**Affiliations:** Axe de Recherche Maladies Infectieuses et Immunitaires, Centre de Recherche du CHU de Québec-Université Laval, Québec, QC, Canada; STELA Dairy Research Center, Institute of Nutrition and Functional Foods, Université Laval, Quebec, Canada

**Keywords:** Milk, transgenic, extracellular vesicles (EVs), microRNAs, *in vitro* digestion, uptake

## Abstract

Milk microRNAs are believed to play gene regulatory functions in the consumer’s cells. Milk from different species is enriched in microRNAs predicted to influence immunity, metabolism, and intestinal homeostasis. For milk microRNAs to regulate gene expression in the consumer, they must survive digestion and be present at sufficient levels to influence intestinal cells and potentially beyond-intestinal cells. Milk microRNAs are proposed to be protected from degradation through their association with milk extracellular vesicles (EVs), which might also deliver them to cells. Studies on milk microRNA oral transfer and tissue bioavailability are limited by interspecies sequence homology, making it difficult to distinguish endogenous from exogenous microRNAs. Here, we used a transgenic (TG) cow model expressing four unique microRNA sequences (AmiRs) in its milk to study their association with milk EVs, their resistance to in vitro digestion, and AmiR uptake and regulatory activity in vitro. We confirmed the presence of the four milk EV populations in raw wild-type (WT) and TG cow milk, similar to those previously reported in commercial (pasteurized) cow milk, and confirmed their association with AmiRs and classical milk microRNAs. AmiRs showed differential resistance to simulated adult digestion. In vitro uptake studies showed a modest gene regulatory effect of AmiRs in Caco-2 cells incubated with TG milk EVs. The intent of using this model was to perform in vitro analysis which could lay the groundwork for later in vivo bioavailability studies, taking advantage of the uniqueness of the AmiRs sequences and bypassing the limitation of microRNA sequence homology.

**Highlights:**

- New milk EV populations identified previously in commercial bovine milk were also identified in raw milk, indicating that they are not merely the result of processing
- The routinely discarded EVs (12K and 35K) seem to be preferentially enriched with microRNAs in raw cow milk as was previously shown for commercial cow milk
- AmiRs in transgenic milk resist differentially to simulated digestion

## Introduction

While food is primarily known for providing nutrients essential for growth, recent research suggests that it may also deliver functional genetic material, with microRNAs being the main focus of such studies. MicroRNAs are small non-coding RNA species (approximately 19-24 nucleotides) that regulate gene expression at the post-transcriptional level and are estimated to regulate up to 60% of human genes ^1,2^. The first study reporting the transfer of dietary microRNAs was by Zhang et al. in 2012, which, despite controversy, reported the transfer of a functional plant microRNA into the tissues and blood of mice upon oral rice intake ^3^. Subsequent studies examined various dietary sources with notable interest in milk microRNAs ^4–6^. MicroRNAs were reported in both raw and processed milk from different species ^7–10^. The profile of milk microRNAs, especially in human, cow, and goat milk, is well-documented, and many of the highly enriched microRNA species are suggested to play critical roles in immunity, intestinal barrier function, and metabolism ^11–19^.

The study of the oral transfer of exogenous microRNAs, including those from milk, is often debated. Among the challenges addressed in this context are those related to the stability of microRNAs during gastrointestinal (GI) digestion, the efficiency of intestinal uptake and tissue distribution, and whether bioavailable levels suffice to alter gene expression. The ability of milk microRNAs to resist digestion is often linked to their association with extracellular vesicles (EVs)-microscopic (50–1,000 nm) structures composed of a cytoplasmic core enclosed by a lipid membrane ^20,21^. EVs mediate material and signal transfer between cells, hence ensuring cell-to-cell communication between neighboring and distant cells ^22–26^. Milk is enriched in EVs with reported roles in immunity and intestinal barrier functions ^27–31^. EVs from milk are enriched in non-coding RNAs, including microRNAs ^7,32–36^. Moreover, newly identified milk EV populations from commercial cow milk (pelleted at 12,000 × g, 35,000 × g, 70,000 × g, and 100,000 × g) with distinct sizes, densities, and protein content, were also shown to present distinct microRNA profiles ^7,33,37^.

Given the association between milk microRNAs and EVs, the latter were suggested to act as vehicles ensuring both the protection and delivery of microRNAs upon oral milk intake. Different studies assessed the resistance of milk microRNAs against simulated digestion using multiple in vitro approaches ^34,38–44^. One study on commercial cow milk reported a 12-26% survivability of microRNAs following simulated digestion under adult conditions ^41^. The authors reported that the majority of these microRNAs were associated with different populations of newly identified EVs ^33,41^. Besides their stability against in vitro digestion, studies also investigated the uptake of milk EVs and/or their microRNA cargo by different cell types in vitro ^45–54^ and assessed their bioavailability and bioactivity in the tissues of milk-consuming subjects ^46,50,54–61^. Nonetheless, besides the intrinsic challenges related to the aforementioned biological barriers, studies assessing the transfer of microRNAs face several technical limitations, like the difficulty in distinguishing exogenous from endogenous microRNAs due to sequence homology across species. To overcome this, several studies employed label-based approaches to label milk EVs and microRNAs, knockout models (lacking one microRNA), or transgenic models with fluorescent EVs to track microRNAs and EVs in cells and tissues ^59,61,62^. However, dye use in complex systems, like in vivo, is limited by issues such as dye detachment or transfer from EVs or microRNAs ^63,64^. Additionally, studies reporting the milk microRNA transfer in vitro and in vivo often lacked data on their biological activity ^55,56^. Thus, the current knowledge on the bioactivity of the transferred microRNAs is limited and rarely quantitatively explored.

In the current study and following up on previous studies in this field, we used milk from a transgenic cow model expressing two unique microRNA sequences (AmiR-3 and AmiR-4) and their star counterparts (AmiR-3* and AmiR-4*) in its mammary gland. This model was originally designed by the AgResearch team in New Zealand to knock down the bovine-specific beta-lactoglobulin (β-lactoglobulin, BLG) protein, which is a major bovine milk allergen ^65–68^. Hence, the intent of this model was to create allergen-free bovine formulas to reduce milk immunoreactivity in infants. Our interest in using the milk from this model, however, was the presence of these unique AmiRs (lacking in mouse and human genomes), which might help in overcoming the sequence homology-associated limitation in microRNA detection. Using milk from this model we set to study the following aspects: 1) the association of AmiRs and other milk microRNAs with the different populations of milk EVs in wild-type (WT) and transgenic (TG) raw cow milk; 2) the digestive fate of TG milk microRNAs following in vitro digestion, 3) the potential of TG milk EVs to transfer microRNAs to cells in vitro and the potential of the latter to regulate gene expression and 4) the minimal copy number of AmiRs required to alter gene expression in recipient cells. Studying these aspects in milk from this transgenic model will set the ground for its use in future in vivo studies to assess the oral transfer of microRNAs while overcoming the limitation associated with sequence homology and hence producing results with minimal bias and enabling more precise tracking and analysis.

## Materials and Methods

### Cow milk samples

Milk samples used in this study were obtained based on a partnership between Dr. Patrick Provost and New Zealand Crown Biotechnology firm AgResearch, including a material transfer agreement (MTA) between Dr. Patrick Provost and Dr. Goetz Laible. The WT milk was obtained from cows that have no direct genetic linkages to the transgenic cows but are of New Zealand dairy genetics, as are the transgenic cows. Samples were collected from cows and then were stored at −80 °C until later use.

### Isolation of milk extracellular vesicles (EVs) by differential ultracentrifugation

Milk EVs were obtained by following a previously described protocol, with slight modifications, that allows quick isolation of milk EVs with very little contaminating proteins ^69^. Raw milk was thawed overnight then centrifuged at 4,000 × g for 30 min to remove cells and fat. Then, the supernatant was subjected to another centrifugation at 4,000 × g for 30 min at 4 °C to allow milk defatting and remove remaining cells and cell debris. One volume of skimmed milk was then mixed with one volume of 2% sodium citrate and kept mixing for 20 min at 4 °C. The Milk:sodium citrate mixture was subjected to centrifugation at 7,200 × g for 10 min to remove sodium citrate-precipitated protein aggregates. Resulting supernatant was then subjected to successive differential ultracentrifugation steps at 12,000 × g (12K) for 2 h, 35,000 × g (35K) for 1 h, then 70,000 × g (70K) for 1 h, and 100,000 × g (100K) for 1 h at 4 °C in a Sorvall WX TL-100 ultracentrifuge, equipped with a T-1250 rotor (both Thermo Fisher Scientific, Waltham, MA, USA). After each step, the pellets were suspended in 1 mL of 0.22 μm filtered sterile phosphate-buffered saline (PBS) pH 7.4.

### Milk EV characterization and EV microRNA quantification

#### Transmission electron microscopy

Isolated EVs were diluted 100X in sodium cacodylate solution (0.1 M, pH=7.3). Samples (10 µl) were then deposited on a grid and left for 3 min to adhere, after which excess liquid was adsorbed with a Kimwipe. Next, 10 µl of uranyl acetate was added to the grid away from light and left to stand for 2 min. Excess uranyl acetate was adsorbed, and the grid was rinsed with distilled water. Excess water was then adsorbed, and grids were left to dry overnight in a desiccator. Images were captured from different areas, and the ones that best represented the overall events observed were chosen. Visualization was made using TEM at the Institut de biologie intégrative et des systèmes, Microscopy Platform at Laval University, Quebec City, QC (80 kV, JEOL® electron microscope 1230, JEOL®, Akishima, Tokyo, Japan).

#### Dynamic light scattering

Zetasizer Nano-ZS dynamic light scattering (DLS) measurement system (Malvern, Worcestershire, UK). The four pellets obtained upon ultracentrifugation were diluted 10X or 100X in 0.22 µm-filtered PBS and deposited in an ultraviolet (UV) cuvette (VWR, Radnor, PA, USA Cat. No. 47743-834). The hydrodynamic size, derived count rate, and polydispersity index were measured at 37 °C with three measurements for each.

#### Protein dosage

Protein dosage for all samples was done using the Pierce BCA Protein Assay Kit (Thermo Fisher Scientific, Waltham, MA, USA, Cat. No. 23227) as per manufacturer’s instructions. Absorbance were read at 562 nm using an EPOCH2 microplate reader (BioTek Instruments, Winooski, VT, USA).

#### Western Blot

Proteins isolated from EVs or from HEK239T cells were mixed with 6X loading buffer (0.15M Tris pH 6.8, 1.2% SDS, 30% glycerol, 1.8% bromophenol blue), heated at 95 °C for 10 min, and subjected to 10% or 12.5% sodium dodecyl sulfate (SDS)-polyacrylamide gel electrophoresis. Proteins were transferred to 0.22 µm (Millipore, Burlington, MA, USA, Cat. No. GVWP00010) or 0.45 µm (Millipore, Burlington, MA, USA, Cat. No. IPVH85R) pore size polyvinylidene fluoride membranes. The membranes were blocked with bovine serum albumin solution (5% bovine serum albumin (BSA) powder milk in tris buffer saline (TBS) with 0.1% Tween 20) for 1 h at room temperature. Membranes were then incubated with primary antibody overnight at 4 °C with monoclonal anti-TSG101 (Abcam, Cambridge, UK, clone 4A10, Cat. No. ab83, diluted 1/500), monoclonal anti-HSP70 (Becton Dickinson (BD), Franklin Lakes, NJ, USA, Cat. No. 554243, diluted 1/1,000), monoclonal anti-ALIX (Santa Cruz Biotechnology, Dallas, TX, USA, clone 3A9, Cat. No. sc-53,538, diluted 1/500), polyclonal anti-CD63 (Santa Cruz Biotechnology, Dallas, TX, USA, Cat. No. sc15363, diluted 1/200), monoclonal anti-XDH (Santa Cruz Biotechnology, Dallas, TX, USA, Cat. No. sc-20991 diluted 1/1,000), polyclonal anti-β-lactoglobulin (Invitrogen, Carlsbad, CA, USA, Cat. No. PA597512, 1/30,000), and monoclonal anti-GAPDH (Abcam, Cambridge, UK, Cat. No. ab9484, diluted 1/2,000). Primary antibodies were diluted in SuperBlock™ Blocking Buffer (Thermo Fisher Scientific, Waltham, MA, USA, Cat. No. 37515). Membranes were then washed three times with TBS tween (3 × 10 min), incubated for 1 h with horseradish peroxidase (HRP)-conjugated secondary anti-mouse (PerkinElmer, Waltham, MA, USA, Cat. No. NEF812001EA, diluted 1/20,000) or anti-rabbit (PerkinElmer, Waltham, MA, USA, Cat. No. NEF822001EA, diluted 1/30,000). Membranes were then washed three times in TBST (3 × 10 min). Western blot signals were revealed with clarity western enhanced chemiluminescence (Bio-Rad Laboratories, Hercules, CA, USA, Cat. No.1705061). The chemiluminescent signal was captured by exposure to the HyBlot CL® Autoradiography 8 x 10", 100 films (Thomas Scientific, Swedesboro, NJ, USA, Cat. No. 1141J52) and developed using the Konica SRX-101A X-ray film processor (Konica Minolta, Tokyo, Japan).

#### RNA extraction from milk samples (skim milk, EVs and supernatant)

RNA extraction was performed as per manufacturer’s instructions with slight modifications. Briefly, 1 ml of RNAzol RT (Sigma-Aldrich, St. Louis, MO, USA, Cat. No. R4533) spiked with 0.15 µl UniSp2 exogenous RNA (QIAGEN, Hilden, NW, Germany, Cat. No. 339390) and added to 250 µl skim milk, milk supernatant (SN), or milk EVs. UniSp2 was used as an exogenous RNA to monitor for variations during RNA extraction. The above mix was added to 400 μL nuclease-free water and mixed by shaking for 20s and then incubated at room temperature for 15 min. Next, the mix was centrifuged at 12,000 × g for 15 min at 4°C, and the supernatant (1,200 µl) was recovered. GlycoBlue (Thermo Fisher Scientific, Waltham, MA, USA, Cat. No. AM9516) and an equal volume of isopropanol were added, mixed gently, incubated for 10 min at room temperature, and centrifuged at 12,000 × g for 10 min at 4 °C. The supernatant was carefully removed, and the pellet was washed twice with 500 µL of 75% ethanol, centrifuging each time at 12,000 × g for 5 min at 4 °C. After air-drying for 10 min, the pellet was resuspended in 12 µL of nuclease-free water and RNA was treated with DNase-I (Invitrogen, Carlsbad, CA, USA, DNA-free™ DNA Removal Kit, Cat. No. AM1906) in accordance with the manufacturer’s protocol.

#### Reverse transcription quantitative PCR (RT-qPCR) of milk samples (EVs, skim milk, and supernatant) RNA

For cDNA generation, miRCURY LNA RT Kit (QIAGEN, Hilden, NW, Germany, Cat. No. 339340) was used on 60 ng RNA of milk samples. UniSp6 RNA, included in the miRCURY LNA RT Kit, was spiked in all RT reactions to control for cDNA synthesis and PCR efficiency. Next, cDNA was diluted 10X and qPCR was performed using miRCURY LNA SYBR Green PCR Kits (QIAGEN, Hilden, NW, Germany, Cat. No. 339345) in 0.1 mL MicroAmp Fast Optical 96-Well Reaction Plate (Applied Biosystems, Cat. No. 4346907) in a StepOne Real-Time PCR instrument (Thermo Fisher Scientific, Waltham, MA, USA) and specific custom LNA oligonucleotides for AmiR-4, AmiR-3*, AmiR-3, AmiR-4* (QIAGEN, Hilden, NW, Germany, Cat. No. 339317, GeneGlobe ID. YCP0055715, YCP005706, YCP0055700, and YCP0055718, respectively), or the LNA PCR primer assays for UniSp2 (QIAGEN, Hilden, NW, Germany, GeneGlobe ID. YP00203950), UniSp6 (QIAGEN, Hilden, NW, Germany, Cat. No. 339306, GeneGlobe ID. YP00203954), miR-148a-3p (QIAGEN, Hilden, NW, Germany, Cat. No. 339306, GeneGlobe ID. YP00205867) and let-7b-5p (QIAGEN, Hilden, NW, Germany, Cat. No. 339306, GeneGlobe ID. YP00204750). We used the following thermal PCR cycle program: denaturation step at 95 [for 2 min, followed by 40 cycles of denaturation at 95 [for 10 s and annealing/elongation at 56 [for 1 min.

#### Standard curve for absolute RNA quantification

The copy numbers of AmiR, miR-148a and let-7b were quantified using a standard curve generated with synthetic RNA oligonucleotides (IDT, Coralville, IA, USA; AmiR-3 synthetic RNA sequence: 5’-AGCUCCUCCACAUACACUCUCGU-3’; AmiR-3* synthetic RNA sequence:5’-CGAGAGUGUGUGGAGGAGCUC-3’ ;AmiR-4 synthetic RNA sequence: 5’-UUUGUAGUCGGUGUCCAGCAC-3’, AmiR-4* synthetic RNA sequence: 5’-CGUGCUGGACCGACUACAAACA-3’, miR-148a synthetic RNA sequence: 5’-UCAGUGCACUACAGAACUUUGU-3’; let-7b synthetic RNA sequence: 5’-UGAGGUAGUAGGUUGUGUGGUU-3’). These oligonucleotides were serially diluted 1:10 to achieve concentrations ranging from 6×10[to 6×10³ copies (per qPCR well), spanning seven orders of magnitude. For each standard curve, the cycle quantification (Cq) values were plotted against the corresponding copy numbers. A linear regression was performed to calculate the equation of the curve and the correlation coefficient (R²). The same was done for the absolute quantification of β-lactoglobulin but the starting material for establishing the standard curve was β-lactoglobulin-coding sequence (represented by cPAEP segment on plasmid map) obtained from β-lactoglobulin plasmid digestion with NdeI (New England Biolabs, Ipswich, MA, USA, Cat. No. R0111S) restriction enzyme (see on plasmid map, **Supplementary Fig. S4**).

### TNO TIM-1 *in vitro* digestion system

#### TIM-1 *in vitro* digestion

TIM-1 is a computer-controlled system which is composed of 4 main digestive compartments mimicking the stomach, duodenum, jejunum, and ileum and are connected through peristaltic valve pumps. Each compartment is made up of glass jacket surrounding a flexible silicone membrane which compresses and relaxes to allow for the peristaltic mixing of the chyme ^70^. The pH, heat and other parameters are continuously monitored through computer-connected sensors. Accordingly, digestive enzymes (e.g., trypsin, pepsin, lipase) and reagents (ex: HCl and bicarbonate) are injected into each compartment to maintain physiologically relevant conditions. For the digestion experiment, transgenic milk (121 ml) was introduced into the system and was digested for 2 h at 37 °C, as this duration approximates the typical residency time of liquid samples. Digestion was performed following a fast-transit protocol as reported by ^71,72^. Parameters of digestion were set to simulate the digestion of a liquid sample in a healthy human adult ^72^ and are listed in the following table:

| *Compartments* | **Compartment residue (initial content) + Secretions** | **Volume (ml)** | **Time to 90% transit completion** | **pH** | **t1/2 / $\beta^*$** |
| --- | --- | --- | --- | --- | --- |
| ***Stomach*** | **Initial content.**<br>• 10 mL of gastric juice (pH=2) 9000 U of pepsin + 375 U of lipase in gastric electrolyte solution<br>**Secretions.**<br>• 900 U/mL of pepsin and 37.5 U/mL of lipase in gastric electrolyte solution (pH=4) circulating at 0.25 mL/min<br>• Water or HCl (1 M) circulating at 0.25 mL/min | 310 | ~ 100 min (calculated based on the exponential formula) | 0/7.4<br>30/3.4<br>60/1.8<br>90/1.5<br>360/1.5 | 30 / 1 |
| ***Duodenum*** | **Initial content.**<br>• 15 g of a pancreatin solution (8%), 30 g of water, 1 mL of trypsin solution (2 mg/mL) and 15 g of small intestine electrolyte solution (pH=7)<br>**Secretions.**<br>• Pancreatin solution (8%) circulating at 0.25 mL/min<br>• Water circulating at 0.50 mL/min<br>• Small intestine electrolyte solution or NaHCO<sub>3</sub> (1 M) circulating at 0.25 mL/min | 55 |  | 6.3 |  |
| ***Jejunum*** | **Initial content.**<br>• Small intestine electrolyte solution<br>**Secretions.**<br>• NaHCO<sub>3</sub> (1 M) if necessary<br>Dialysis fluid:<br>• Small intestine electrolyte solution circulating at 10 mL/min | 115 |  | 6.5 |  |
| *Ileum* | **Initial content.**<br>• Small intestine electrolyte solution<br>**Secretions.**<br>• NaHCO<sub>3</sub> (1 M) if necessary<br>Dialysis fluid:<br>• Small intestine electrolyte solution circulating at 10 mL/min | 115 | ~ 311 min<br>(calculated based on the exponential formula) | 7.4 | 160/1.8 |

\*Gastric and ileal deliveries are modelled with a power exponential formula: $f = 1 - 2^{-(t/t_{1/2})^\beta}$ , where f represents the fraction of meal delivered, t the time of delivery, $t^{1/2}$ the half-time of delivery, and $\beta$ is the coefficient describing the shape of the curve.

#### TIM-1 Media

Freshly prepared working solutions contained pepsin derived from porcine gastric mucosa (Sigma-Aldrich, St. Louis, MO, USA, P6887), trypsin sourced from porcine pancreas (Sigma-Aldrich, St. Louis, MO, USA, T9201), pancreatin extracted from porcine pancreas (Sigma-Aldrich, St. Louis, MO, USA, 4 × USP, P1750), and lipase from Rhizopus oryzae (Amano Enzyme USA Co., Ltd., Elgin, IL, USA, LIPASE DF-DS), all dissolved in sterile deionized water. The electrolyte solutions consisted of a gastric electrolyte mixture (6.2 g/L NaCl, 2.2 g/L KCl, 0.3 g/L CaCl[, and 1.5 g/L NaHCO[) and a small intestinal electrolyte solution (5.0 g/L NaCl, 0.6 g/L KCl, and 0.3 g/L CaCl[) ^73,74^.

#### TIM-1 sample collection

Sampling was done from each of the four TIM-1 compartments at 0, 30, 60, and 120 min of digestion. At each time point, the effluent was collected, and its volume was noted. Samples were flash-frozen in liquid nitrogen and stored at −80 °C for later RT-qPCR analysis.

#### RNA extraction and RT-qPCR of TIM-1 samples

RNA extraction was performed using TRIzol-LS reagent (Invitrogen, Carlsbad, CA, USA, Cat. No. 10296028) followed by DNase-I treatment (Invitrogen, Carlsbad, CA, USA, DNA-free DNA Removal Kit, Cat. No. AM1906) in accordance with the manufacturer’s protocol. An exogenous synthetic control microRNA (Caenorhabditis elegans let-7-as mutant, 2 fmol; IDT, Coralville, IA, USA) was added during the TRIzol-LS homogenization step to monitor for variations during RNA extraction. HiFlex miScript RTII kit (QIAGEN, Hilden, NW, Germany, Cat. No. 218160) was used for reverse transcription. cDNA was generated from 1,000 ng of HEK293T RNA. Next, cDNA was diluted 10X and qPCR was performed using SSo Advanced SYBR Green mix (Bio-Rad Laboratories, Hercules, CA, USA, Cat. No. 1725721) in 0.1 mL MicroAmp Fast Optical 96-Well Reaction Plate (Applied Biosystem, Foster City, CA, USA, Cat. No. 4346907) in the StepOne Real-Time PCR System (Applied Biosystem, Foster City, CA, USA, Cat. No. 4376357) and the custom-designed primers with proprietary sequences were used for the four AmiRs (QIAGEN, Hilden, NW, Germany, Cat. No. 339317).

### Caco-2 transfections and incubation with EVs

#### Caco-2 cells

Caco-2 cells were obtained from American type culture collection (ATCC) and were grown in complete Dulbecco’s modified Eagle’s medium (DMEM, Wisent Inc., Saint-Jean-Baptiste, QC, Canada, Cat. No. 319-005-CL) supplemented with 10% decomplemented (56 °C, 30 min with occasional shaking) fetal bovine serum (Wisent Inc., Saint-Jean-Baptiste, QC, Canada, Cat. No. 080-150), 100 Units/ml of penicillin, 100 µg/ml streptomycin and 2mM L-glutamine (Wisent Inc., Saint-Jean-Baptiste, QC, Canada, Cat. No. 405-202-EL). Cells were grown and maintained in tissue culture flask (Thermo Fisher Scientific, Waltham, MA, USA, Cat. No. 13-680-65) and incubated at 37 °C in a humidified atmosphere under 5% CO2. Cells were kept in the exponential growth phase and subcultured every 4-5 days.

#### Plasmid constructs

To generate the plasmid constructs, synthetic DNA inserts, containing the AmiR-4 binding site (WT BS) or a mutant (Mut. BS) AmiR-4 binding site (flanked with restriction sites of Not-I and Xho-I) were designed (IDT, Coralville, IA, USA) to be cloned into the 3′ untranslated region (3′UTR) of the *Renilla luciferase* (Rluc) gene within the psiCHECK-2 (Promega, Madison, WI, USA, Cat. No. C8021) dual-luciferase reporter plasmid. The plasmid has an ampicillin resistance gene to allow for bacterial selection during plasmid cloning. The plasmid also expresses a *firefly luciferase* (FLuc) gene under a separate promoter, which serves as an internal control for normalization of transfection efficiency and reporter activity. Cloning was performed using the restriction enzyme digestion and ligation approach. Both the psiCHECK-2 backbone and the synthetic DNA insert were digested with NotI (NotI-HF® (New England Biolabs, Ipswich, MA, USA, Cat. No. R3198S) and XhoI (New England Biolabs, Ipswich, MA, USA, Cat. No. R0146S) which cleaved at sequences flanking the Rluc 3′UTR region in the plasmid and AmiR-4 binding site in the insert to create cohesive ends. To enhance ligation efficiency, promote correct insert orientation, and prevent self-ligation, the digested insert was subsequently treated with calf intestinal alkaline phosphatase (New England Biolabs, Ipswich, MA, USA, Cat. No. M0525S) to remove 5′ phosphate groups. Following dephosphorylation, the insert was ligated into the digested vector backbone using T4 DNA ligase (New England Biolabs, Ipswich, MA, USA, Cat. No. M0202S). The ligation product was transformed into *E.coli* DH5α competent cells, and positive clones were selected on LB agar plates containing 100 µg/mL ampicillin (Wisent Inc., Saint-Jean-Baptiste, QC, Canada, Cat. No. 400-110-XG). Plasmid DNA was extracted from individual colonies using a miniprep kit (Bio-Basic, Markham, ON, Canada, EZ-10 Spin Column Plasmid DNA Miniprep Kit, Cat. No. BS414), and the presence and correct orientation of the insert were confirmed by DNA sequencing at the Genomics Center (Genomics Center at the Research Center of Laval University, Quebec, QC, Canada).

#### Cell transfection and dual luciferase assay

For the transfection experiment, 50,000 Caco-2 cells were cultured in a 6-well plate and transfected 48 h later with psiCHECK-2 plasmids (described above, 50 ng per well) using Lipofectamine™ 2000 (Thermo Fisher Scientific, Waltham, MA, USA, Cat. No. 11668019). Six hours following plasmid transfection, the medium was removed and replaced with medium containing 60 μg TG EVs or 60 μg WT EVs (mix of all EV fractions). Twenty-four hours following EV addition, cells were washed with PBS and lysed using Dual-Luciferase™ Reporter (DLR™) Assay Systems (Promega, Madison, WI, USA, Cat. No. E1980). The RLuc and FLuc bioluminescence signals were measured using the Dual Glo reagents of the DLR reported assay following the manufacturer’s instructions. Light emission was measured using a luminometer (TECAN INFINITE M1000 PRO, Tecan Austria GmbH, Grödig, Salzburg, Austria). Rluc activity was expressed relative to the expression of the Fluc internal control. RLuc expression was further normalized to the control in which cells were transfected with the plasmid with and incubated with medium without EVs. All assays were conducted in triplicate in a 96-well format.

### HEK293T transfections

#### HEK293T

HEK293T cells were obtained from American type culture collection (ATCC) and were grown in complete Dulbecco’s modified Eagle’s medium (DMEM, Wisent Inc., Saint-Jean-Baptiste, QC, Canada, Cat. No. 319-005-CL) supplemented with 10% decomplemented (56 °C, 30 min with occasional shaking) fetal bovine serum (Wisent Inc., Saint-Jean-Baptiste, QC, Canada, Cat. No. 080-150), 100 Units/ml of penicillin, 100 µg/ml streptomycin and 2mM L-glutamine (Wisent Inc., Saint-Jean-Baptiste, QC, Canada, Cat. No. 405-202-EL). Cells were grown and maintained in a tissue culture flask (Therma Fisher Scientific, Waltham, MA, USA, Cat. No. 13-680-65) and incubated at 37 °C in a humidified atmosphere under 5% CO2. Cells were kept in the exponential growth phase and subcultured every 2–3 days.

#### HEK29T cell plasmid and microRNA mimic transfection

The β-lactoglobulin plasmid used in the transfection experiments was custom-designed and purchased at VectorBuilder Inc. (VectorBuilder, Chicago, IL, USA, Vector ID: VB900139-1033mxb, see plasmid map in **Supplementary Fig. S4**). For the transfection experiment, 38,000 cells were cultured in a 24-well plate and transfected the following day using the calcium/phosphate method with 50 ng plasmid per well. Twelve hours following plasmid transfection, cells were washed with PBS and transfected with the custom AmiR-3 and AmiR-4 microRNA mimics (Horizon Discovery, Cambridge, UK, AmiR-3 Cat. No. CTM-1043164; AmiR-4 Cat. No. CTM-1043164) or Cel-miR-67 mimic negative control (confirmed to have minimal sequence identity with miRNAs in human, mouse, and rat; Horizon Discovery, Cambridge, UK, CN-001000-01-05) using Lipofectamine™ RNAiMAX (Thermo Fisher Scientific, Waltham, MA, USA, Cat. No. 13778150) as per manufacturer’s protocol. The plasmid, the mimics as well as the transfection reagents were diluted in Opti-MEM (Invitrogen, Carlsbad, CA, USA, Cat. no. 31985062). Twenty-four hours following mimic transfection, cells were washed with PBS, collected by trypsin detachment and directly added to 1X RIPA lysis buffer (150 mM NaCl, 1% Nonidet-P-40 (NP-40), 0.5% sodium deoxycholate, 0.1% SDS, 50 mM Tris, pH 7.4) supplemented with protease inhibitors (cOmplete EDTA-free, Roche, Basel, Switzerland, Cat. No. 4693132001; and PhosSTOP, Roche, Basel, Switzerland, Cat. No. 4906845001) for protein isolation or washed twice with PBS and then added to RNAzol RT (Sigma-Aldrich, St. Louis, MO, USA, Cat. No. R4533) reagent for RNA extraction.

#### RNA extraction and RT-qPCR of HEK293T samples

Total RNA was isolated from HEK293T cells using RNAzol RT (Sigma-Aldrich, St. Louis, MO, USA, Cat. No. R4533) according to the manufacturer’s guidelines. The RNA samples were treated with DNase I (Invitrogen, Carlsbad, CA, USA, DNA-free™ DNA Removal Kit Cat. No. AM1906) and quantified using a BioDrop-μLITE Spectrophotometer (Isogen Life Science, De Meern, Netherlands). HiFlex miScript RTII kit (QIAGEN, Hilden, NW, Germany, Cat. No. 218160) was used for reverse transcription. cDNA was generated from 300-400 ng of HEK293T RNA. Next, cDNA was diluted 10X and qPCR was performed using the SSo Advanced SYBR Green mix (Bio-Rad Laboratories, Hercules, CA, USA, Cat. No. 1725721) in 0.1 mL MicroAmp^TM^ Fast Optical 96-Well Reaction Plate (Applied Biosystems, Foster City, CA, USA, Cat. No. 4346907) in a StepOnePlus Real-Time PCR System (Thermo Fisher Scientific, Waltham, MA, USA) and the following custom-designed primers (IDT, Coralville, IA, USA) GGGGACTTGGTACTCCTTGG (forward) and CACACTCACCGTTCTCCCAT (reverse) for β-lactoglobulin (amplicon size 152bp); and ACCACAGTCCATGCCATCAC (forward) and TCCACCACCCTGTTGCTGTA (reverse) for GAPDH (amplicon size 452bp).

#### Western Blot

Proteins isolated (10-15 μg) HEK239T cells were mixed with 6X loading buffer (0.15M Tris pH 6.8, 1.2% SDS, 30% glycerol, 1.8% bromophenol blue), heated at 95 °C for 10 min, and subjected to 10% or 12.5% sodium dodecyl sulfate (SDS)-polyacrylamide gel electrophoresis. Proteins were transferred to 0.22 µm (Millipore, Burlington, MA, USA, Cat. No. GVWP00010) or 0.45 µm (Millipore, Burlington, MA, USA, Cat. No. IPVH85R) pore size polyvinylidene fluoride membranes. The membranes were blocked with skim milk solution (5% powder milk in tris buffer saline (TBS) with 0.1% Tween 20) or bovine serum albumin (5% powder BSA in TBS with 0.1% Tween 20) for 1 h at room temperature. Membranes were then incubated with primary antibody overnight at 4 °C with polyclonal anti-β-lactoglobulin (Invitrogen, Carlsbad, CA, USA, Cat. No. PA597512) and monoclonal anti-GAPDH (Abcam, Cambridge, UK, Cat. No. ab9484). Membranes were then washed three times with TBS tween (3 × 10 min), incubated for 1 h with horseradish peroxidase (HRP)-conjugated secondary anti-mouse (PerkinElmer, Waltham, MA, USA, Cat. No. NEF812001EA) or anti-rabbit (PerkinElmer, Waltham, MA, USA, Cat. No. NEF822001EA). Membranes were then washed three times in TBST (3 × 10 min). Western blot signals were revealed with clarity western enhanced chemiluminescence (Bio-Rad Laboratories, Hercules, CA, USA, Cat. No.1705061).

#### Statistical analysis and test

We clarify that all transgenic milk samples but not wild-type samples, were obtained from the same transgenic animal, collected at three widely separated time points; as these represent repeated measures rather than independent biological replicates, we acknowledge this limitation while noting that generating a larger cohort of transgenic animals was not feasible for this study.

All statistical analyses were performed using Prism 10.4.1 (GraphPad Software Inc., San Diego, CA, USA). Data are presented as mean ± standard deviation (SD) unless otherwise stated. Due to the small sample size (n = 3 per group), normality could not be formally assessed. Therefore, for comparisons between two groups, two-tailed unpaired t-tests were used to compare WT and TG samples (for the comparison of microRNA copy number between WT and TG milk in milk and EV populations, and for comparison of protein content, diameter, and derived count rate between WT and TG milk). P-values were adjusted for multiple testing using the Benjamini–Hochberg false discovery rate procedure; FDR-adjusted p-values <0.05 were considered statistically significant used. For comparisons involving more than two groups (comparison between the protein and mRNA levels between the conditions with different mimic concentrations), a one-way analysis of variance (ANOVA) was performed followed by Tukey’s post hoc test for multiple comparisons. Statistical significance was set at p < 0.05. All experiments, except TIM-1 *in vitro* digestion experiments, were conducted with at least three independent biological replicates (n ≥ 3), and each condition included technical replicates to ensure reproducibility.

## Results

### Detection of AmiRs in transgenic (TG) milk

Given that microRNAs in milk originate principally from mammary gland epithelial cells ^75,76^, we assumed that AmiRs would also be secreted into the milk from TG cows. Milk samples were first collected from different TG and WT cows or from the same cow at different time points and then defatted to obtain skim milk. Using RT-qPCR, AmiRs copy number was assessed in TG skim milk samples as well as in WT skim milk as a control. Given that miR-148a and let-7b are among the top ten microRNAs consistently reported to be highly enriched in milk, we included the analysis of their levels in our samples ^32,75,77^. Absolute quantification of copy number revealed that AmiR-3 was the most highly enriched relative to the other three AmiRs **(Figure 1B)**. Interestingly, the induced expression of the four AmiRs (via transgenesis) resulted in substantially high levels of the AmiRs in comparison with miR-148a and let-7b **(Figure 2B-C)**. Notably, sRNA sequencing revealed that transgenesis affected the expression of certain classical milk microRNAs, altering their relative levels in TG versus WT milk, despite not being direct targets **(Supplementary Fig. S1).** For instance, miR-99a-5p reads were reduced tenfold in transgenic milk compared to wild-type milk, while let-7f reads were four times higher in transgenic milk compared to those in wild-type milk **(Supplementary Fig. S1)**. Overall, these results confirm the secretion of the four AmiRs into TG milk.

**Figure 1.**
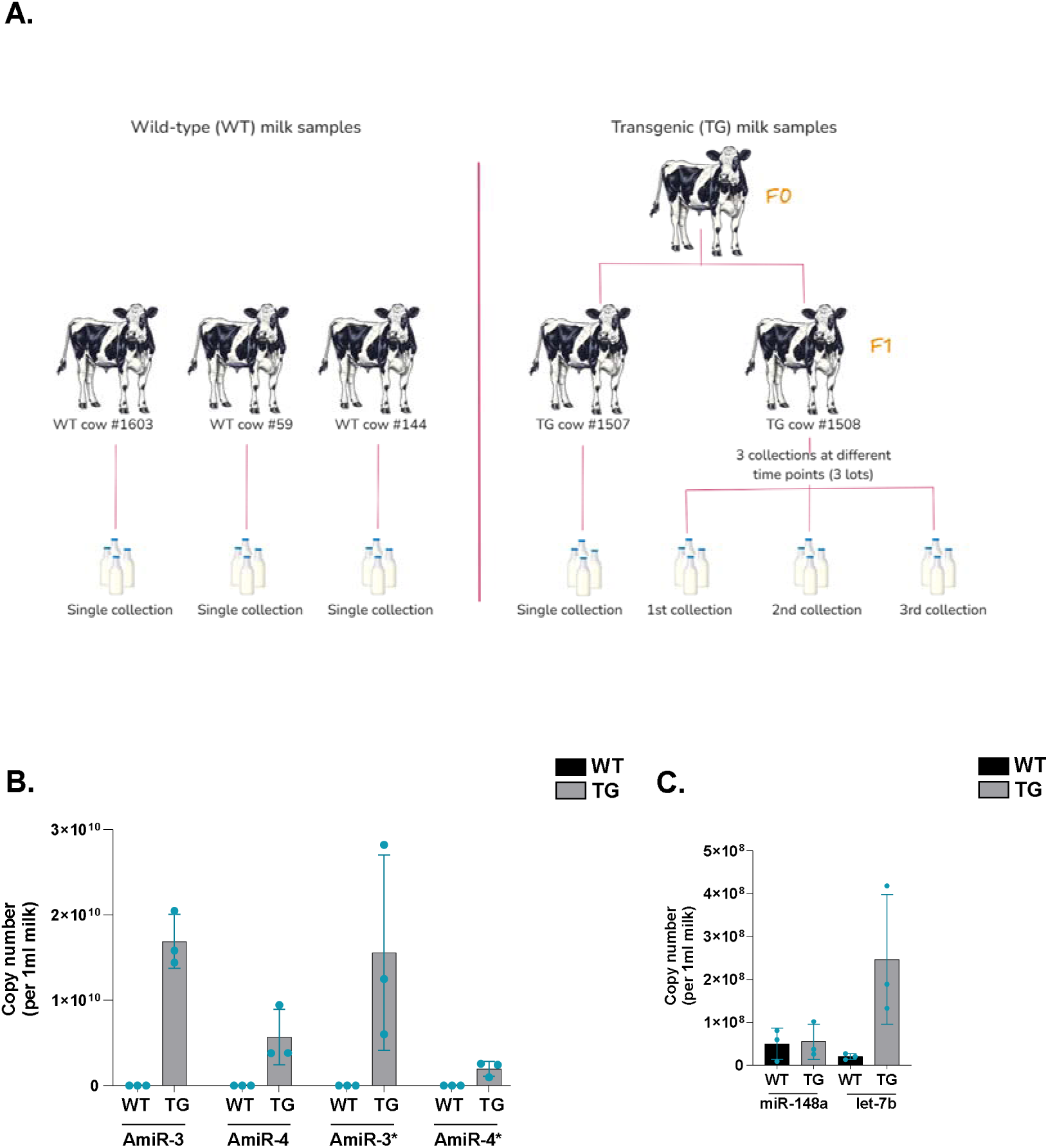
The transgenic (TG) and wild-type (WT) milk samples used in this study and quantification of their microRNAs. **A.** Collection of TG and WT milk samples. The TG milk used in this study was collected from two F1 generation transgenic cows. One lot was collected from each of the three different WT cows (#1603, #144, and #59) while three lots were collected from TG cow #1508 at three different time points. **B.** Real-time qPCR-based absolute quantification of AmiR-3, AmiR-4, AmiR-3*, AmiR-4*, and the two classical milk microRNAs, miR-148a and let-7b, in WT and TG non-fractioned skim milk. Copy number is presented per 1ml milk EVs of non-fractioned skim milk. UniSp2 RNA spike-in was used to normalize the RT-qPCR data. The values are means ± SD of three independent samples representing 3 lots of each milk type (n=3). Unpaired t-test was used to compare WT and TG samples; p-values were adjusted for multiple comparisons using the Benjamini–Hochberg procedure.

**Figure 2.**
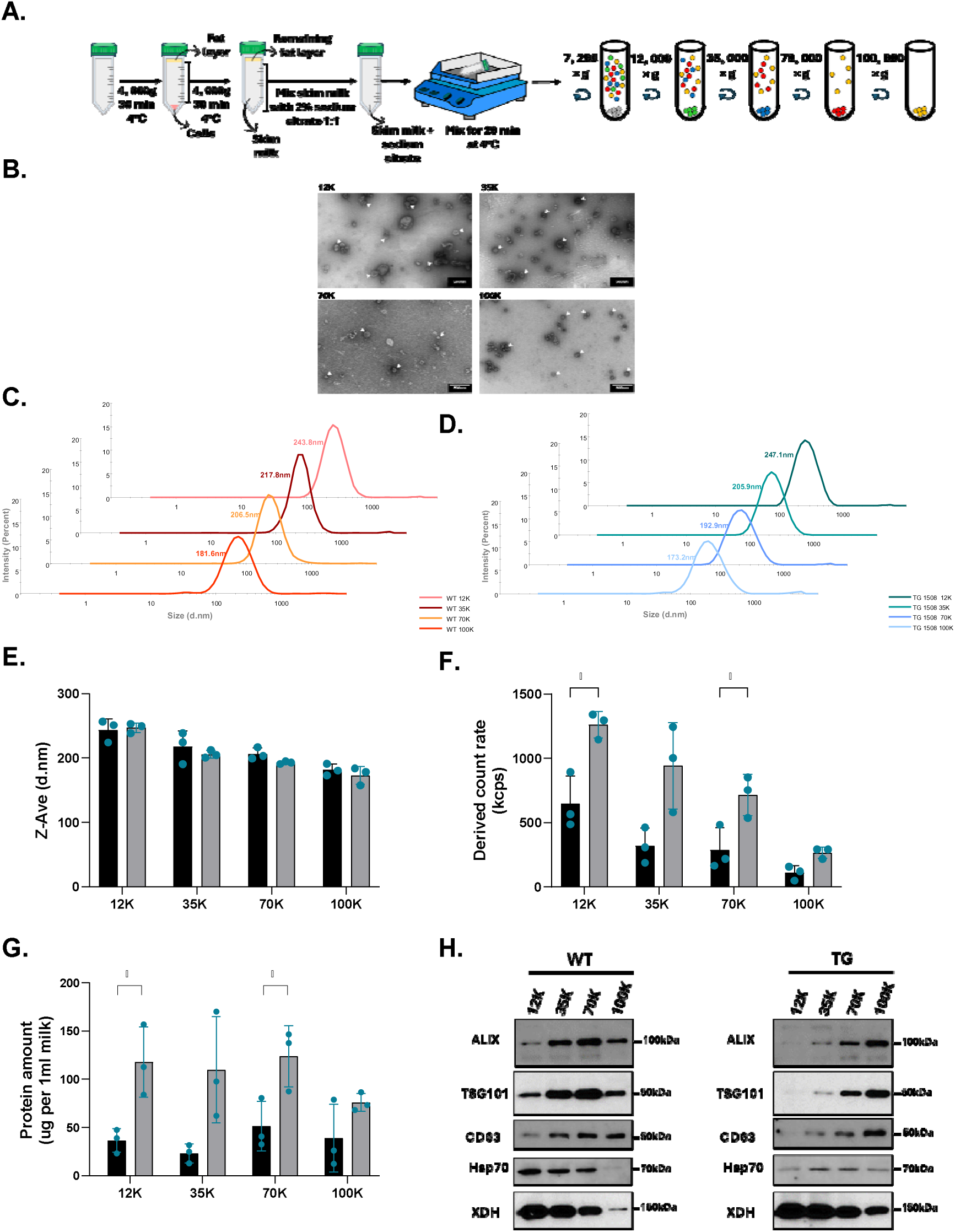
Isolation and characterization of WT and TG raw milk EV populations. **A.** Scheme depicting the workflow of milk de-cellularization, defatting, and sodium citrate treatment followed by EV isolation via differential ultracentrifugation. **B.** Transmission electron microscopy images of the four pellets obtained upon differential ultracentrifugation of TG skim milk. Raw peaks showing the size distribution by intensity of particles in each of the four 12K, 25K, 70K, and 100K populations (presented are the averaged peaks of the three milk lots (biological triplicates) with three readings recorded per sample (technical triplicates) in **C.** WT and **D.** TG milk. **E.** Measurements of EV hydrodynamic size (Z-average) using dynamic light scattering (DLS). Shown are the Z-average measurements of the four EV populations isolated from three TG (TG #1508 lot 1, lot 3 and lot 3) and three WT (WT #1603, #59 and #144) skim milk lots (n=3) **F.** DLS estimate of the particle count (scattering intensity) in each of the four EV populations of TG and WT milk lots (n=3). **G.** Quantification of the protein content (in µg) in each EV subset obtained from 1ml milk or from 1ml non-fractioned skim milk or SN (n=3). The values are means ± SD of three independent experiments (n=3). For each condition (e.g., 12K, 35K, etc.) unpaired t-test was used to compare WT and TG samples; p-values were adjusted for multiple comparisons using the Benjamini–Hochberg procedure (*adjusted p<0.05). **H.** Western blot of milk EV canonical protein markers.

### Isolation and characterization of EV populations from raw WT and TG milk

Processed (commercial) cow milk was proven to contain four different populations of EVs with distinct physicochemical properties, protein, and microRNA content. Using differential ultracentrifugation, these EVs could be pelleted at 12, 000 × g, 35, 000 × g, 70, 000 × g, and 100, 000 × g and are hence designated as 12K, 35K, 70K, and 100K EVs (EV pellets), respectively. To determine whether similar populations exist in raw cow milk, we applied the same ultracentrifugation protocol described in ^69^ to isolate these four EV populations from skim WT and TG milk **(Figure 2A)**. To study the physical and chemical properties of our EV isolations, we used dynamic light scattering (DLS), transmission electron microscopy (TEM), and western blot **(Figure 2)**.

Images from TEM showed the typical cup-shaped morphology of TG EVs and a rather heterogeneous in terms of size among different pellets, as well as within the same pellet (**Figure 2B and Supplementary Fig. S2A** for average diameter calculated with Image J).

DLS measures the hydrodynamic size of particles inferred from how the particles scatter the light, without providing visuals of their morphology. The standard deviation of particle diameter (Z-average) was consistently narrow across the three biological triplicates for both WT and TG pellets as shown in **Figure 2C-D**. Comparative analysis of the hydrodynamic size (Z-average) showed no significant variation between matching EV subsets from WT and TG milk (e.g. 12K WT vs 12K TG, **Figure 2E and Supplementary Fig. S2B)**. Notably, all TG pellets showed higher particle count (derived count rate estimates particle count in a suspension) compared to their corresponding WT pellets (**Figure 2F and Supplementary Fig. S2C)**. Consistent with the higher particle count, protein yield in all TG pellets was higher than in WT pellets, with a significant increase in the 12K and 70K populations **(Figure 2G)**.

Protein extracts from the four pellets of both WT and TG milk were probed with antibodies against proteins commonly known to exist in EVs (either as surface or cytosolic proteins). Common EV-associated markers like ALIX, TSG101, CD63, and Hsp70 (all of which are involved in various EV biogenesis pathways) were detected in both WT and TG pellets though at varying levels **(Figure 2H)**. Other proteins like Xanthine dehydrogenase (or XDH which is known as marker for milk fat globules but was also found associated with milk sEVs (“exosomes”) ^78^) were also detected in all WT and TG pellets. Overall, we were able to detect the common EV markers in both WT and TG pellets.

### Partitioning and distribution of AmiRs and classical milk microRNAs over the four EV populations and supernatant (SN)

To study the association of AmiRs with the EVs, RNA was extracted from the four pellets and the absolute levels of the four AmiRs (per 1 ml milk) were calculated using RT-qPCR. The four AmiRs, AmiR-3, AmiR-3*, AmiR-4, and AmiR-4*, were the most enriched in the 12K pellet, while their levels progressively decreased with increasing centrifugal force **(Figure 3A-D)**. A similar trend was also seen for the two classical milk microRNAs, miR-148a and let-7b in both WT and TG milk **(Figure 3E-F)**. All four AmiRs, as well as miR-148a and let-7b, were least enriched in the supernatant (SN) obtained from the last centrifugation at 100K.

**Figure 3.**
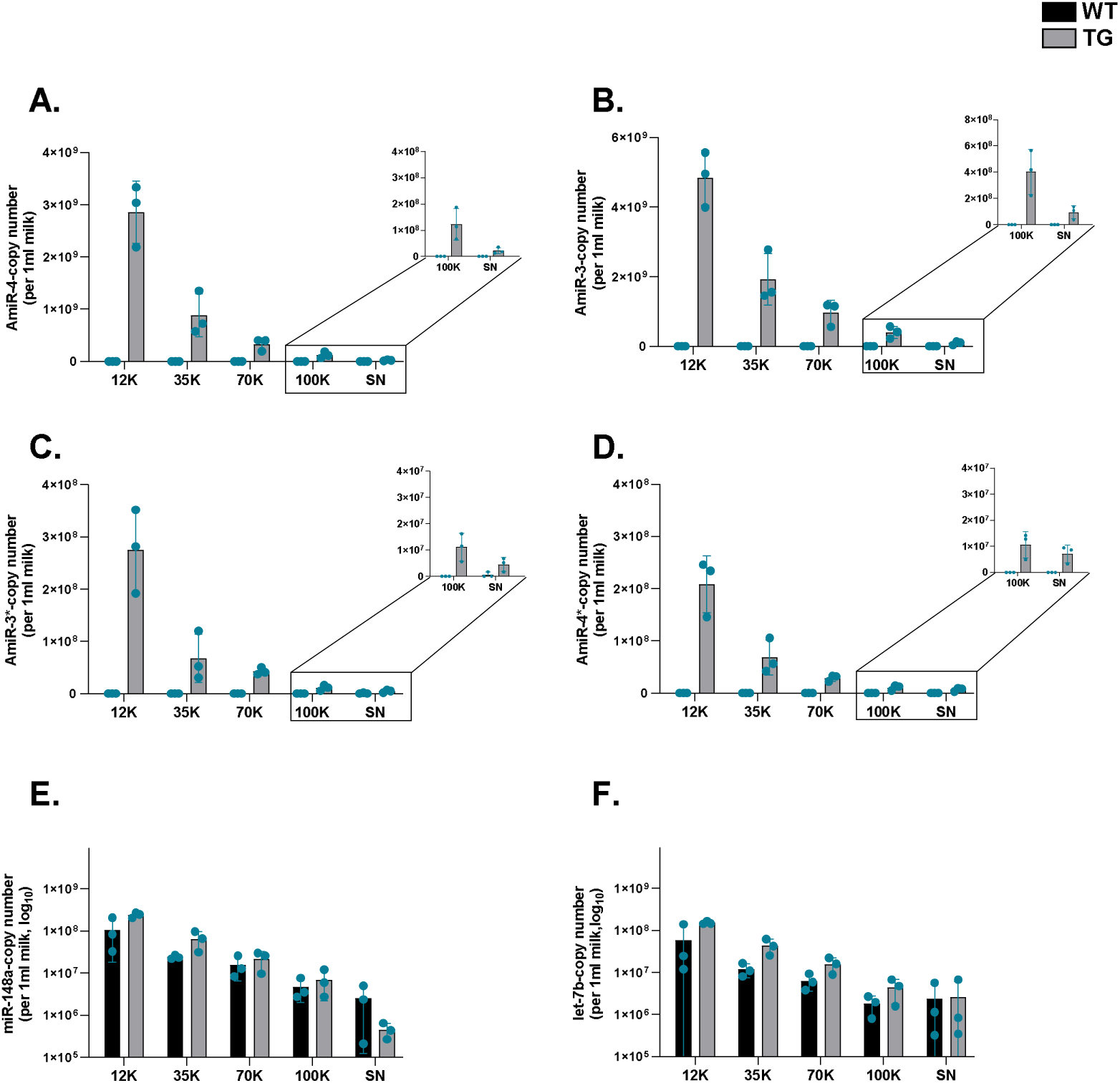
Quantification of the microRNAs in EV subsets of WT and TG milk. Real-time qPCR-based absolute quantification of **A.** AmiR-3, **B.** AmiR-4, **C.** AmiR-3*, **D.** AmiR-4*, and the two classical milk microRNAs, **E.** miR-148a and **F.** let-7b, in the 4 EV subsets and SN of WT and TG milk, as well as in non-fractioned skim milk. Copy number is presented per 1ml milk EVs, 1ml SN, or 1ml non-fractioned skim milk. UniSp2 and UniSp6 RNA spike-ins were used to normalize the RT-qPCR data. Reported errors are the standard deviation of three values from three independent samples representing the three lots (WT #1603, #44, #59, and TG #1508 lot 1, 2,3) of each milk type. Unpaired t-test was used to compare WT and TG samples; p-values were adjusted for multiple comparisons using the Benjamini–Hochberg procedure.

### *In vitro* simulated digestion of transgenic milk

Given that EVs are viewed as vehicles that can protect their microRNA content against harsh conditions (like digestion) and that AmiRs were found to majorly associate with TG-milk EVs, we decided to assess the ability of AmiRs to resist to *in vitro* digestion. Here, we tested the survivability of AmiRs against *in vitro* simulated digestion using the TNO TIM-1 system ^70^ **(Figure 4A and Supplementary Fig. S3).** We simulated healthy adult digestion parameters over two hours using the protocol from ^41^. After two hours (2 h), we collected digests from each compartment as well as from the effluent (digests that has passed through all system compartments). We also collected effluent samples at T=0.5 h and T=1 h. We used RT-qPCR to quantify the copy number of AmiRs in the total volume of each compartment. AmiRs seem to be differentially partitioned over the different TIM-1 compartments after 2 h of digestion **(Figure 4B-E)**.

**Figure 4.**
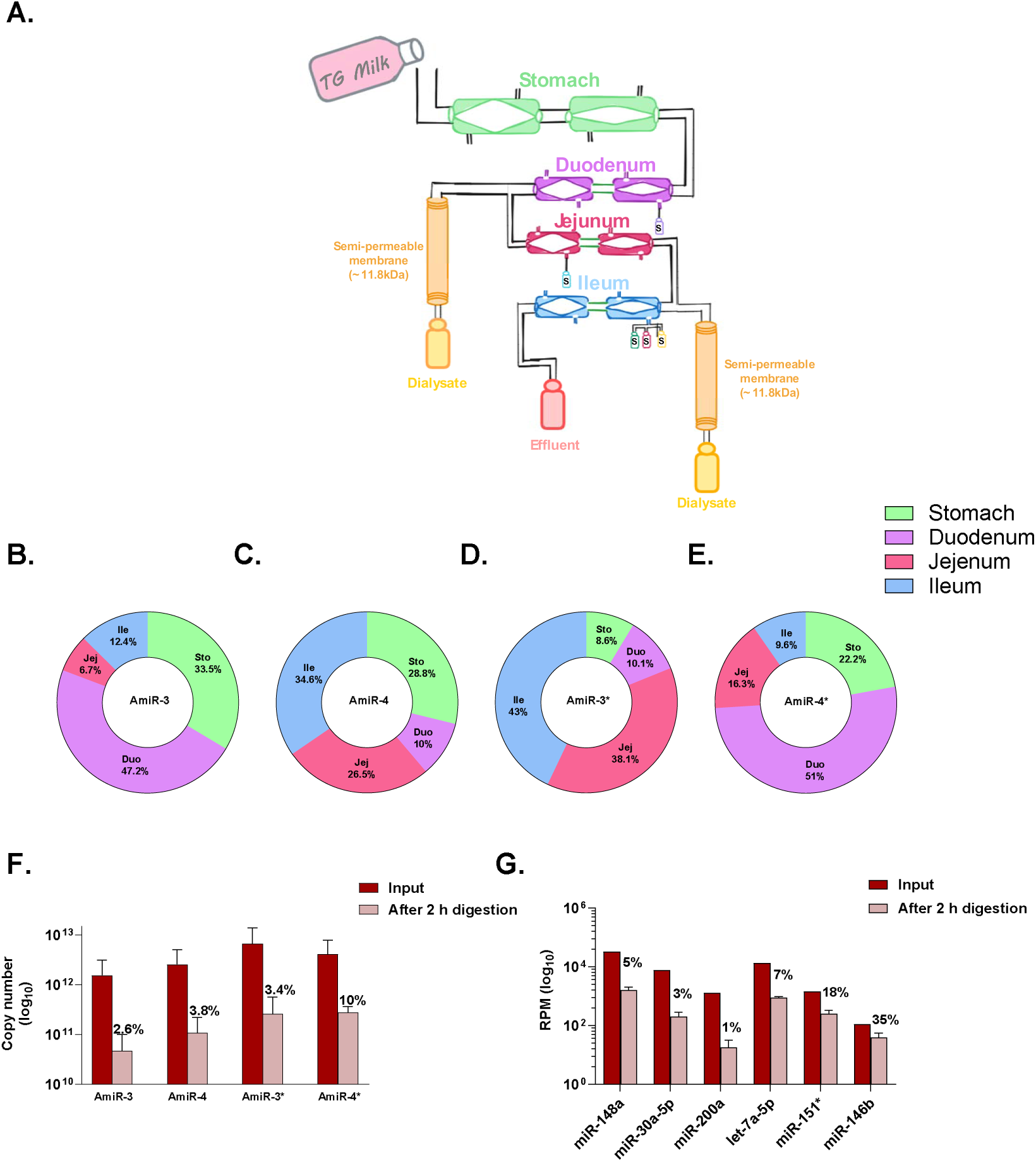
*In vitro* simulated digestion of transgenic milk. Defatted transgenic (TG) milk was subject to *in vitro* simulated digestion using the TNO TIM-1 system. **A.** Scheme showing the different parts of TIM-1. Donut charts showing the percentage (percentage of total microRNA in all compartments) of **B.** AmiR-3, **C.** AmiR-4, **D.** AmiR-3*, and **E.** AmiR-4* in each of the four TIM-1 compartments: stomach, duodenum, ileum, and jejunum following 2-h digestion. **F.** The number of copies of each of the four AmiRs was calculated in the effluents that had exited the system at T0.5 h, T1 h, and T2 h. Effluents are the samples that have traversed the entire system and were collected at the peristaltic valve located at the ileum. Presented is the sum of the copy number in the total volume of the effluent collected at each time point (Eff0.5 h + Eff1 h + Eff2 h). Percentages represent the quantity of microRNAs remaining relative to the initial input. The data are displayed as mean ± SD, n=2. **G.** Small RNA-seq data showing the reads per million of six classical milk microRNAs before (input) and after 2-h digestion. Percentages represent the quantity of microRNAs remaining relative to the initial input. The data are displayed as mean ± SD.

Figure 4F shows the AmiRs that were detected in the effluents collected at T0.5 h, T1 h and T2 h (summed up). These AmiRs have passed through all system compartments and, therefore, withstood the entire digestion process extending over 2 h. This graph indicates the differential resistance of AmiRs against digestion with AmiR-4* resisting the most, as 10% of its initial input remained detectable after exiting the system as effluent. AmiR-4 and AmiR-3* showed similar survivability (around 3.5% remaining) while AmiR-3 resisted the least with more than 97.3% lost (Figure 4F). Classical milk microRNAs showed varying resistance to simulated digestion (Figure 4G). miR-146b retained 35% of miR-146b of its initial input, while only 1% of miR-299a persisted after 2 h (Figure 4G). Though the majority of AmiRs seem to be lost during digestion, still however, in terms of copy number, more than 10^12 copies of each AmiR resisted digestion and were still detectable by RT-qPCR (Figure 4G).

### Assessing the possibility of TG EV-mediated transfer of AmiRs into cells in vitro and AmiRs’ potential gene-regulatory effects

In order for the microRNAs in milk to exert gene regulatory functions, they should be transferred into the cells where they could bind to their target mRNA. Given that few cellular models endogenously express the target of AmiRs (β-lactoglobulin), we designed a plasmid in which we inserted the binding site (mutated (Mut. BS) or not (WT BS)) of AmiR-4 in the 3’UTR of RLuc in psiCHECK-2 plasmid (Figure 5A). The binding of AmiR-4 to its WT BS in the mRNA of RLuc but not to Mut. BS will reduce RLuc protein expression which will be reflected on its bioluminescence. Compared to control (no EVs), TG EVs were able to induce a decrease in the RLuc signal in cells transfected with WT BS plasmid (Figure 5B). Nonetheless, this decrease was not significant when compared to RLuc signal in TG EV-treated cells which were transfected with Mut. BS plasmid. Cells incubated with WT EVs, transfected with either plasmid, did not show significant change in RLuc signal when compared to the control (no EVs) (Figure 5B).

**Figure 5.**
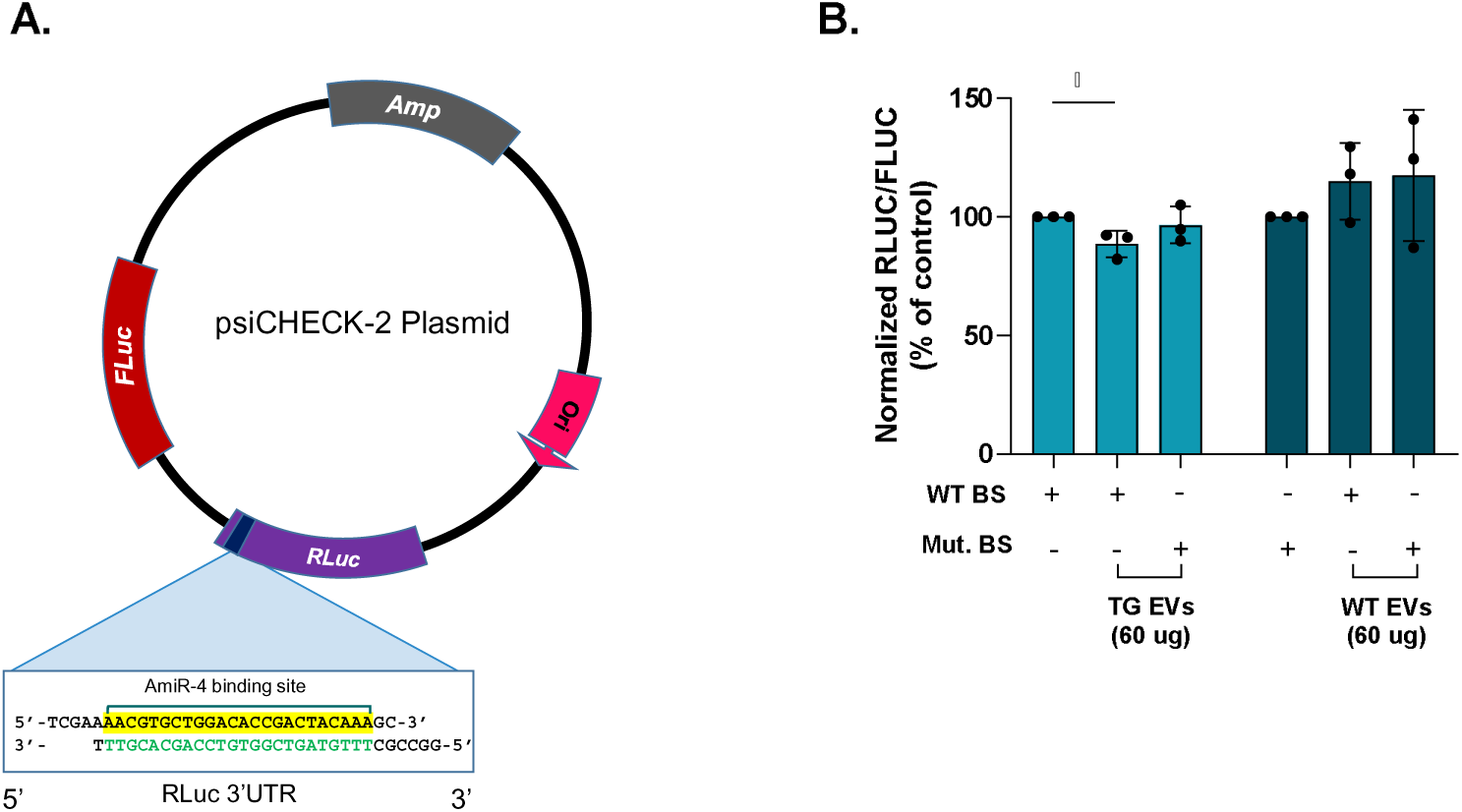
Assessing the potential of TG EVs to deliver AmiR-4 into cells in vitro and to induce, if any, gene-regulatory effects. Caco-2 cells were transfected with the psiCHECK-2 plasmid carrying the AmiR-4 binding site or AmiR-4 mutant binding site inserted in the 3’UTR of the RLuc gene. **A.** Scheme depicting plasmid design and highlighting the position of the AmiR-4 binding site (mutant or not). **B.** Luciferase bioluminescent signal of RLuc was normalized to that of FLuc and expressed as a percentage of control (no EVs). Data are presented as mean ± SD (n=3). Statistical significance was assessed using an unpaired *t*-test for pairwise comparisons between individual groups (*p<0.05). Each pair of conditions was compared independently. The absence of a significance symbol indicates no statistically significant difference. WT BS: plasmid carrying the exact binding site sequence of AmiR-4. Mut. BS: plasmid carrying the mutated binding site sequence of AmiR-4; 3’UTR = 3’ untranslated region; RLuc = Renilla luciferase; FLuc = Firefly luciferase; Ori = Origin of replication; Amp = Ampicillin.

### Estimation of the “effective” copy number of AmiRs and equivalent EV amount

Given that the experiment with the psiCHECK-2 plasmid did not provide conclusive results, we decided to use another plasmid construct carrying the β*-lactoglobulin* gene which could better recapitulate the interaction between AmiR-4 and its target, especially that the binding sites of AmiRs in the transgenic cow are found within the open reading frame of β*-lactoglobulin* gene. Moreover, given that the observed reduction in RLuc signal was relatively modest, we decided to estimate the “effective” copy number of AmiRs required to alter gene expression *in vitro* which could better help estimate the amount of EVs needed. For that, we transfected HEK293T cells with β-lactoglobulin-expressing plasmid (see **Figure 6A and Supplementary Fig. S4-5** for plasmid map and AmiR binding sites in β*-lactoglobulin* gene).

**Figure 6.**
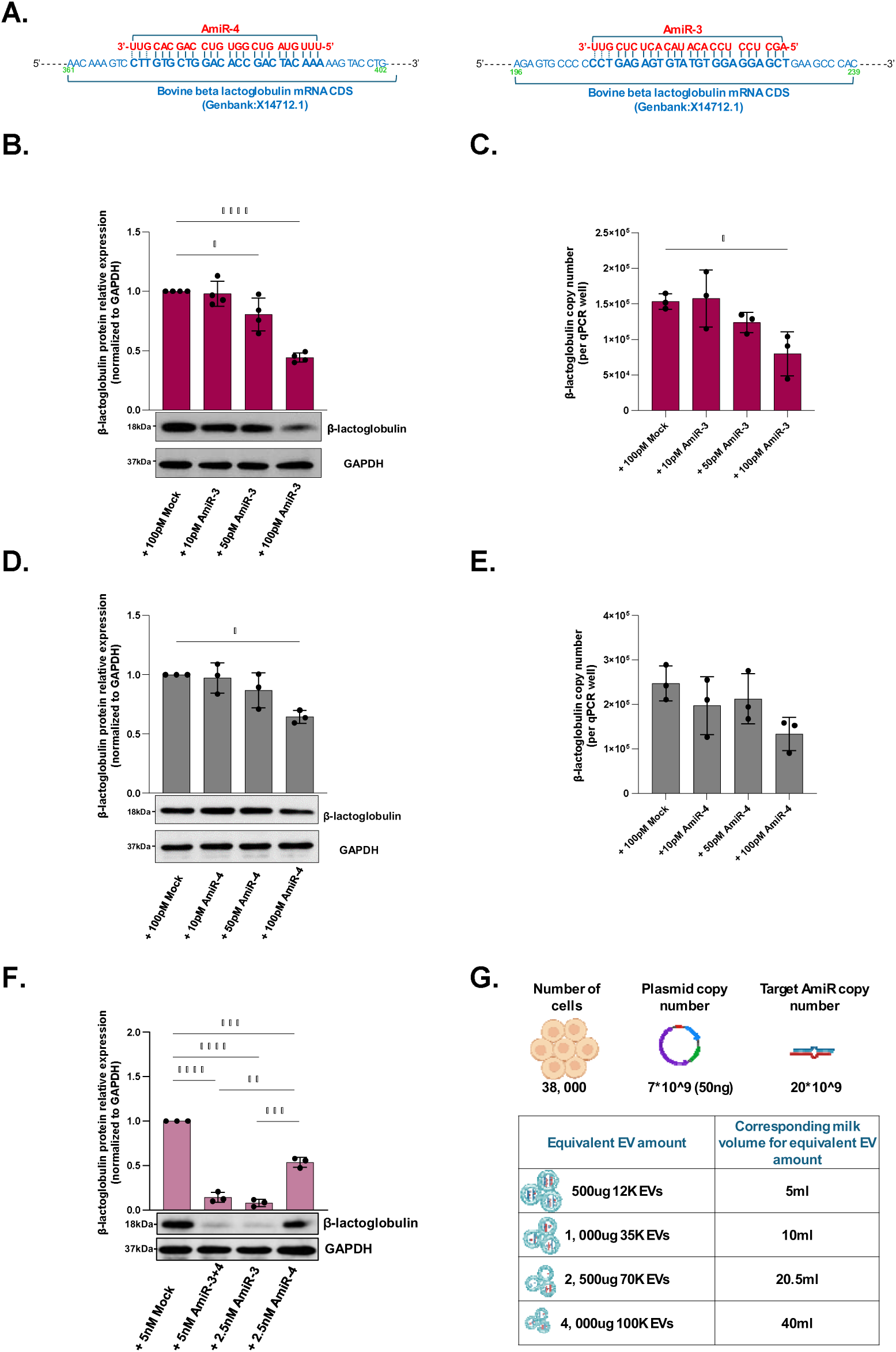
Estimation of the AmiR “effective” copy number. HEK293T cells were co-transfected with a plasmid expressing β-lactoglobulin along with different concentrations of AmiR-3/4 mimics. **A.** Scheme depicting the binding sites of AmiR-3 and AmiR-4 in the coding sequence of β-lactoglobulin mRNA (Genbank no. X 14712.1). **B-C** Western blots, densitometry, and RT-qPCR-based absolute quantification of β-lactoglobulin protein and mRNA levels in HEK293T cells after co-transfection with β-lactoglobulin plasmid and AmiR-3 mimic at three different concentrations (10 pM, 50 pM, and 100 pM). **D-E** Western blots, densitometry, and RT-qPCR-based absolute quantification of β-lactoglobulin protein and mRNA levels in HEK293T cells after co-transfection with β-lactoglobulin plasmid and AmiR-4 mimic at three different concentrations (10 pM, 50 pM, and 100 pM). **F.** Western blots and densitometry quantification of β-lactoglobulin protein levels in HEK293T cells after co-transfection with β-lactoglobulin plasmid, AmiR-3, and AmiR-4 (simultaneously or individually). The microRNA mimic mock, Cel-67, was used as a negative control at 100 pM or 5 nM. Proteins and total RNA were extracted from cells and used for analysis 24 h following mimic transfection. GAPDH was used as an internal control for western blot densitometry analysis and for RT-qPCR. The values are means ± SD of three independent experiments (n=3-4). Statistical analysis was performed using one-way ANOVA followed by Tukey’s post hoc test (*p<0.05, **p<0.01, ***p<0.001, ****p<0.0001). **G.** Table showing the equivalent copy number of AmiRs required to inhibit β-lactoglobulin protein and/or mRNA expression under specific experimental conditions (transfection of 38,000 HEK293T cells with 50ng β-lactoglobulin-expressing plasmid). CDS = Coding sequence.

Since plasmids are often built with strong promoters and often result in overexpression of the gene of interest, we tried to use a minimum plasmid amount to match the expression levels of endogenous genes, which will hence require less AmiRs to influence gene expression. We tested several AmiR-to-plasmid ratios and their effect on β-lactoglobulin protein and mRNA levels **(Supplementary Fig. S6)**. AmiR-3 significantly reduced β-lactoglobulin protein at both 100 pM and 50 pM but only decreased mRNA levels at 100 pM (Figure 6B-C). In contrast, AmiR-4 inhibited protein expression only at 100 pM, with no significant effect at the other concentrations or at the β-lactoglobulin mRNA level (Figure 6D-E).

In their paper, Jabed *et al.* mentioned that AmiR-4 and AmiR-3 have a synergetic effect. Hence, we also tested any potential synergetic effect of the two AmiRs at 5 nM (2.5 nM AmiR-3 + 2.5 nM AmiR-4) while comparing it with the standalone effect of each AmiR. Figure 6F shows that AmiR-3 alone reduced β-lactoglobulin protein to lower levels than AmiR-3 and AmiR-4 combined. Hence, AmiR-3 and AmiR-4 do not seem to act synergistically upon the transfection of their synthetic sequences into HEK293T cells.

A simple estimation was made to determine the amount of EVs (in µg) necessary to deliver the effective AmiR copy number (∼10^9 copies, ∼2 × 10^4 copies per cell) under our experimental conditions (Figure 6G). This analysis showed that 100 µg, 500 µg, 2,500 µg, and 4,000 µg of the 12K, 35K, 70K, and 100K EV fractions, respectively, are needed to reach the target copy number.

## Discussion

Research on the transfer of dietary microRNAs as intact, functional molecules capable of modulating gene expression in consumers’ cells remains controversial ^6,79–81^. The controversy stems from both technical limitations and intrinsic biological barriers, which could influence the detection, stability, and amount of the transferred microRNAs, and eventually their gene regulatory functions ^79^. One technical limitation is represented by the high degree of sequence homology of microRNAs across species, making it challenging to track the dietary microRNAs in the recipient’s cells or tissues. Furthermore, the complexity of the food matrix makes it difficult to causally link increased microRNA levels to ingested sources. Moreover, the use of label-based approaches may yield false positives due to non-specific label transfer or detachment from microRNAs or EVs[^55,63,64^. As for the biological barriers, orally ingested microRNAs must overcome degradation in the GI tract, mucus entrapment, stool excretion, and dilution across tissues.

One way to address the limitation emerging from sequence homology in the study of the oral transfer of dietary microRNAs would be to use a dietary source containing unique microRNA sequences lacking in animal models (e.g., mice). As a basis for such future in vivo studies, we studied milk EVs as delivery and protective vehicles of microRNAs in milk from a transgenic cow model expressing two unique microRNA sequences (AmiR-3 and AmiR-4) and their star counterparts (AmiR-3* and AmiR-4*). We investigated milk EV populations in WT and TG milk, their AmiR and microRNA contents, AmiR resistance to in vitro digestion, and the capacity of TG EVs to transfer AmiRs to cells in vitro.

The detection of intact microRNA molecules in both raw and processed bovine milk products indicates an unusual stability of these molecules, which are otherwise highly vulnerable to degradation ^7,9–11,76,82,83^. The documented stability of extracellular microRNAs against physical and chemical (including enzymatic) treatments is often attributed to their association with EVs ^41,84,85^. Milk from different sources is enriched in EVs carrying a wide spectrum of RNAs, lipids, and proteins with potential biological functions ^86–90^. Benmoussa *et al.* and others reported new populations of EVs (12K, 35K, and 70K) in pasteurized commercial cow milk, which presented different chemical and physical properties compared to the commonly studied milk small EVs (sEVs, or 100K EVs; also referred to as “exosomes”) ^7,37,91–93^. Furthermore, both the 12K and 35K populations were proven to carry the bulk of milk total microRNAs ^33^. In the current study, we reported that the four EV populations (12K, 35K, 70K, and 100K) are also present in raw, non-treated cow WT and TG milk (Figure 2). The characterization of these EV populations revealed similar properties (size, protein markers) as those reported in commercial cow milk ^33^ (Figure 2). This suggests that these EV populations likely do not originate from milk fat globules during homogenization and are not a product of processing. Additionally, similar to microRNAs in commercial milk ^33^, classical milk microRNAs (in WT and TG milk) as well as AmiRs (in TG milk) were majorly associated with EVs compared to the supernatant (SN, EV-depleted milk fraction), and more enriched in the 12K and 35K compared to the 70K and 100K (Figure 3). We believe, however, that using milk samples from different animals, rather than multiple samples from the same animal as it is the case in this study, would be necessary to increase the statistical power and confirm such enrichment. This partitioning of microRNAs is critical, especially since these larger EV populations (12K and 35K) are often discarded during standard milk EV isolation pipelines ^47,51,53,55^. This finding underscores the need to revisit EV isolation strategies to fully capture the wide spectrum of milk EVs.

The resistance of milk EVs and microRNAs to digestion was previously assessed using several models of in vitro digestion. Studies using models based on a simple mixing of milk with digestive enzymes revealed an overall stability of bovine and human milk microRNAs to simulated digestion ^38,39,44^. Yet another report, where a similar digestion approach was used, indicated a high overall degradation of milk microRNAs following digestion ^54^. In the current study, the dynamic, compartmentalized TIM-1 digestion system was employed to simulate adult digestion using the protocol by Benmoussa *et al.* ^41,70,94^. In their study on the effect of digestion on commercial (pasteurized) bovine milk microRNAs, the authors showed that 12% of bta-miR-223-3p and 26% of bta-miR-125b-5p sustained a two-hour digestion process ^41^. The aforementioned study reported that the most degradation of both microRNAs (miR-223-3p and bta-125b-5p) occurred in the stomach ^41^. However, others indicated that the intestinal digestive conditions caused a higher loss of microRNAs compared to the gastric digestion ^54,61^. In the current study involving the digestion of TG raw cow milk, the four AmiRs partitioned differently over the four TIM-1 compartments, which might reflect differential microRNA stability (Figure 4B-E). This could result from AmiR-specific characteristics influencing their behavior under the varying pH and enzymatic conditions of each compartment. Additionally, microRNA ratios, which have sustained the whole digestion process (in the effluent), also indicate a differential survivability for both the AmiRs and the other classical milk microRNAs (Figure 4F-G). Similar differential resistance has been reported in other studies and is suggested to be related to factors like microRNA secondary structure, primary sequence, CG content, and associated protein partners ^38,41,42^. Although much of the AmiRs were lost during digestion, the remaining copy number was still considerable. This is noteworthy as under normal dietary patterns (or under typical feeding conditions), surviving microRNAs could accumulate with repeated milk intake. Moreover, even if microRNA levels are too low for systemic distribution, they may still affect intestinal cells locally. This highlights the potential effect of milk microRNAs on intestinal homeostasis in intestinal diseases and during development ^18,95–97^. The resistance of milk microRNAs to in vitro digestion was also corroborated by studies on bovine-specific milk microRNAs detected in the stool of formula-fed human infants ^58^.

Following oral milk intake in an in vivo setting, milk components, including milk EVs, will first encounter intestinal cells following digestion. Surviving EVs might establish contact hubs with intestinal cells, leading to the internalization and the release of their microRNA cargo, which could then regulate gene expression by binding target mRNAs. In the current study, the incubation of Caco-2 cells transfected with a plasmid carrying the AmiR-4 binding site (downstream of the RLuc gene) with TG EVs resulted in only a slight reduction in luciferase expression, which was not significant when compared to the cells transfected with a plasmid carrying the mutant AmiR-4 binding site (Figure 5B). Because the experiment yielded non-significant results, we cannot infer a gene-regulatory role for microRNAs or their transfer into cells by EVs. Accordingly, the next section examines possible factors that could have affected these outcomes.

Starting with the AmiR copy number, to perform their typical gene regulatory functions, a minimum microRNA copy number per cell is required ^98^. Here, we found that a minimum of ∼10^9 copies (equivalent to ∼2 × 10^4 per cell) of AmiR-3 or AmiR-4 mimics need to be transfected into cells to induce a significant reduction in β-lactoglobulin protein levels. This copy number is equivalent to AmiRs in 100 µg, 500 µg, 2,500 µg, and 4,000 µg of 12K, 35K, 70K, and 100K EVs, respectively (under the given experimental conditions, (Figure 6G)). Based on this estimation, the amount of EVs (60 µg) used with Caco-2 might not be sufficient for effective gene regulation, which might explain the modest reduction reported here. Also, regarding the physiological relevance, having enough levels of AmiRs while factoring in the 90% or so digestion-based loss and the number of cells in the intestinal barrier (if considering a local effect of the ingested milk microRNAs) in vivo (Figure 4F), might require the ingestion of supraphysiological levels of milk in a single intake. Nonetheless, this number of transfected mimics can not be directly extrapolated to an equivalent amount of EVs due to the difference in the uptake efficiencies between EVs and transfection reagents ^99^. The uptake of EVs by the cells might not be as efficient. Also, many EVs might fail to escape endolysosomal degradation, delivering their cargo to the degradative lysosomes rather than to the cytosol ^100^. Recent findings suggest that only 24% of internalized EVs may successfully deliver their cargo to the cytoplasm and that this occurs in only about 5% of recipient cells ^101,102^. Furthermore, in studies on milk EV uptake, it was reported that milk EVs are internalized by cells via caveolin-mediated endocytosis among other endocytic pathways ^50,52,53,103^. However, this pathway might be weakly active in Caco-2 cells, as these do not express caveolin-1 ^104^, a key limitation of this experiment. This could thus lower the EV uptake efficiency in these cells, hence the importance of testing other intestinal cell lines in the future. Furthermore, though, spatially speaking, intestinal cells are likely the immediate target of milk EVs, the latter might rather have a large influence on the immune cells of the gut-associated lymphoid tissue (GALT) or cells of the intestinal microbiota. The primordial effects of 35K and 100K EVs of commercial cow milk on a model of colitis were at the level of immune responses and immune factors ^91^. It is thus important to include immune cells in the study of milk EV-cell interactions. Besides factors related to recipient/target cells, the pipeline used for EV isolation might have influenced EV integrity or their surface proteins, key factors in EV-cell and EV-extracellular matrix (ECM) interactions ^105^. It could also be hypothesized that, as shown for some classical microRNAs when comparing WT and TG milk (**Supplementary Figure S2**), the transgenesis process could have influenced the expression of other genes, like EV surface proteins or other proteins related to their uptake. This highlights one potential limitation of using this model or other models with genetically-modified EVs ^56,59,61^.

We acknowledge several limitations in the current study. First, the use of multiple milk samples derived from the same animal rather than from different animals, the use of Caco-2 cells which are deficient in one of the endocytic pathways, and the data of digestion being replicated only twice (n = 2). Although TIM-1 provides continuous, real-time control of digestion parameters and duplicate runs are commonly used ^106,107^, additional independent digestion replicates, increased numbers of animals, and in vivo validation of microRNA resistance to digestion will be required to confirm the bioaccessibility of milk microRNAs.

## Conclusion

The demonstrated resistance of milk AmiRs to digestion and their association with EVs support the use of TG milk as a model dietary source for studying inter-species oral microRNA transfer in vivo, helping to bypass sequence homology-related limitations. However, potential alterations in EV protein composition due to transgenesis (e.g., β-lactoglobulin, a bovine milk EV protein[^37,108^) should be considered when interpreting the results, as they may limit extrapolation to wild-type milk EV biodistribution.

## Author contribution

Z.H. conceived and designed the study. Z.H., A.B., and N.M. performed the experiments and collected the data (A.B and N.M. TIM-1 experiment, Z.H. all other experiments). I.F. provided expertise, guidance, advice, and analytical tools (TIM-1 experiment). Z.H. and A.B. analyzed the data. Z.H. wrote the manuscript. All authors read and approved of the final manuscript.

## Acknowledgements

We thank Dr. Patrick Provost for his major role in this project, including designing the study, supervising the project, designing key experiments, and providing the funding support. This work was supported by the Canadian Institutes of Health Research (CIHR) (SIRUL: 123293, to Dr. Patrick Provost). We also thank Dr. Goetz Laible for providing the milk samples, without which this project would not have been possible. We also extend our sincere thanks to Dr. Caroline Gilbert for the invaluable guidance in supervising the writing process of this paper and for her assistance with the figures and data analysis. We extend our gratitude to Dr. Idrissa Diallo and Dr. Marine Lambert for their guidance, advice, and helpful discussions throughout the course of this work.

## Data availability statement

Data supporting the findings of this study, involving the RNA-seq data, are available from the corresponding author upon reasonable request.

## Supplementary figures

**Supplementary Fig. S1.**
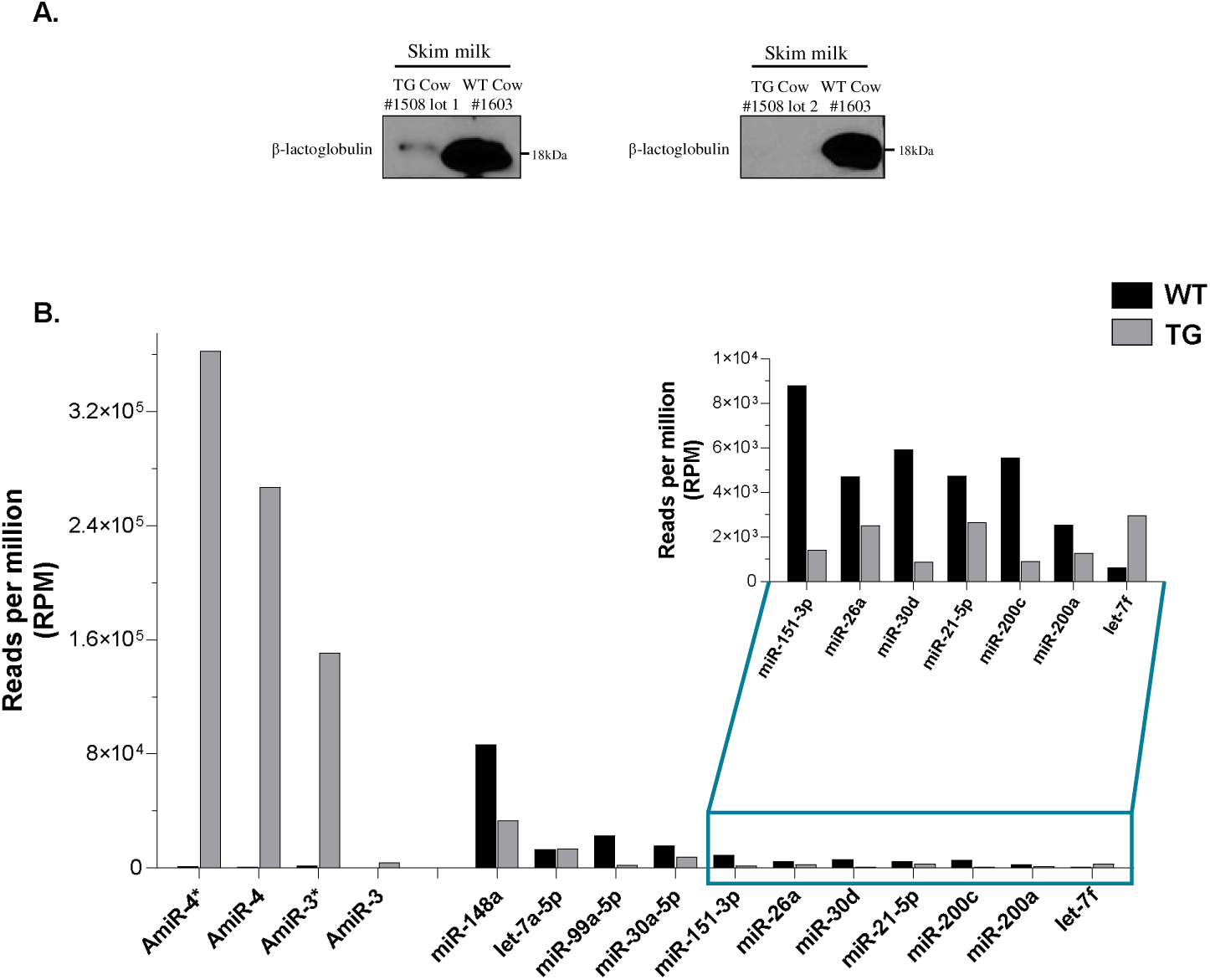
**A.** Detection of β-lactoglobulin protein in WT (WT cow #1603 lot) and TG (TG #1508 lot 1 and lot 2) non-fractioned skim milk samples. **B.** Small RNA sequencing of TG (TG #1507) and WT (TG # 1603) milk. Presented are the reads per million (RPM) of the four AmiRs (AmiR-4*, AmiR-4, AmiR-3*, and AmiR-3) along with 11 other classical milk microRNAs. These classical milk microRNAs are commonly known to be highly enriched in milk of different types and from different species.

**Supplementary Fig. S2.**
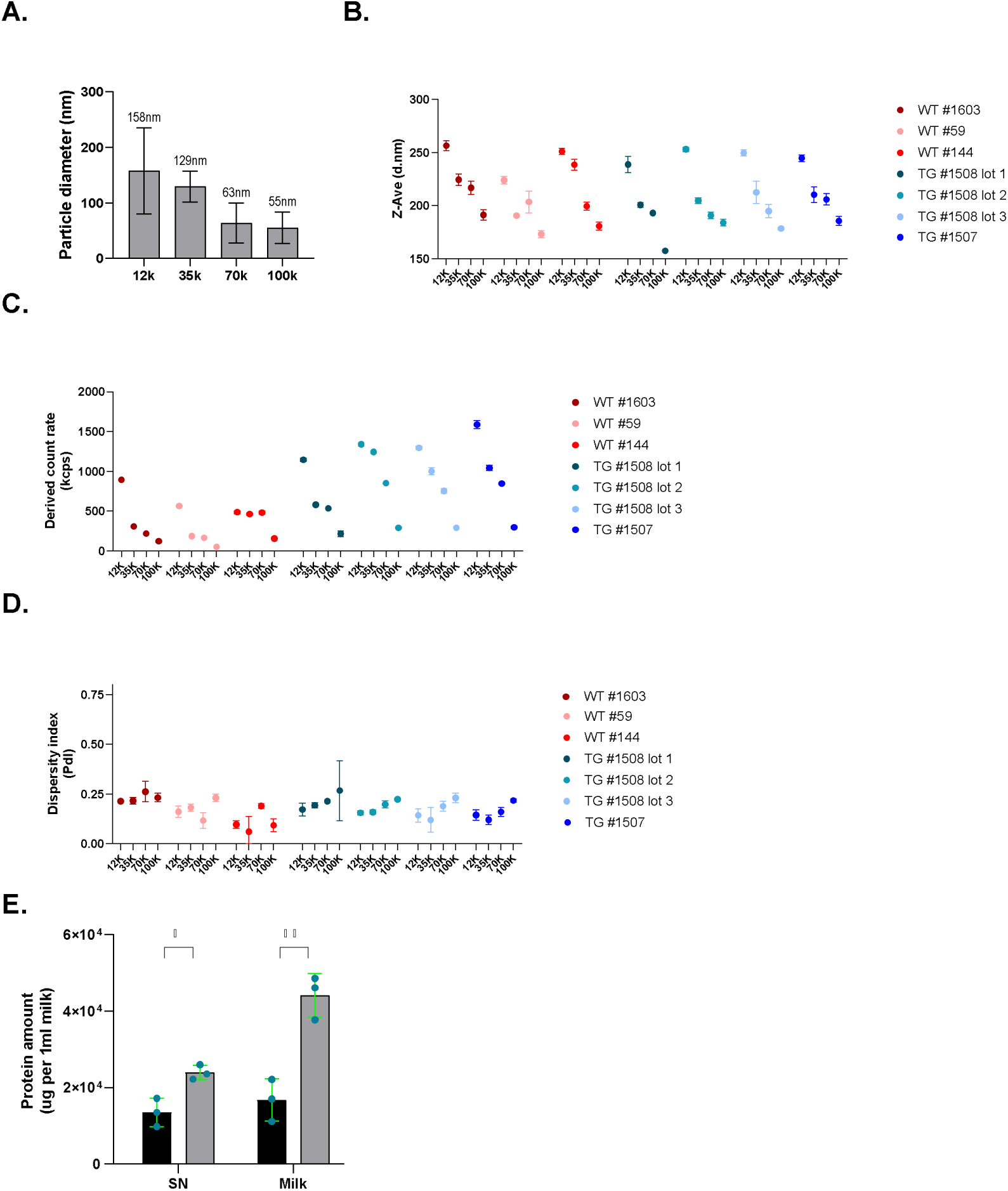
Characterization of WT and TG raw milk EVs. **A.** Transmission electron microscope images of the four TG EV subsets (12K, 35K, 70K, 100K); magnification 18,500x; scale=500 nm. **A.** Measurements of EV diameter using ImageJ software tools on TEM images. The values are means ± SD of measurements from different photos. **B.** Measurements of EV hydrodynamic size (Z-average) using dynamic light scattering (DLS). The values are means ± SD of three independent experiments (n=3, 3 lots of milk). **C.** DLS estimate of the particle count (scattering intensity) in each of the four EV subsets of TG and WT milk lots (n=3). The values are means ± SD of three independent experiments (n=3). An unpaired t-test was used as a statistical test. **D.** Dispersity index measured by DLS as an estimate of the size uniformity of particles in different pellet samples (n=3). Shown are the measurements of the four EV subsets isolated from four TG (TG #1508 lot 1, lot 2, and lot 3, and TG #1507) and three WT (WT #1603, #59, and #144) skim milk lots, reported errors are the standard deviation of three measurements. **E.** Quantification of the protein content (in µg) in 1ml non-fractioned skim milk or SN (n=3). The values are means ± SD of three independent experiments (n=3). For each condition (e.g., SN, milk) an unpaired t-test was used to compare WT and TG groups (*p<0.05, **p<0.01).

**Supplementary Fig. S3.**
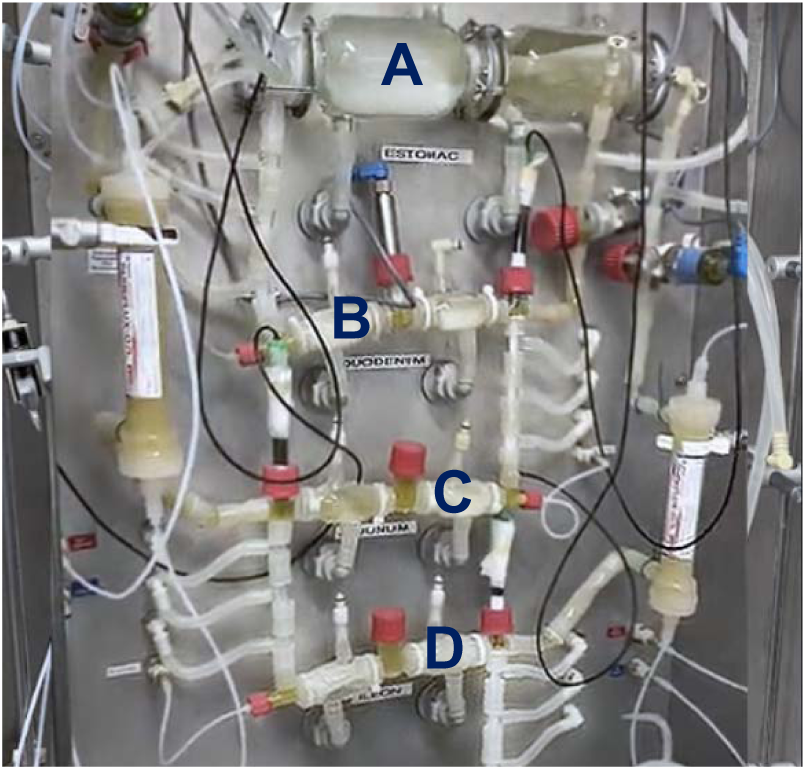
TNO TIM-1 in vitro digestion system. Photo of the system present at the STELA Dairy Research Center, Institute of Nutrition and Functional Foods (INAF), Université Laval, Quebec, Canada. TIM-1 consists of 4 main compartments connected through peristaltic valve pumps, representing the stomach (A), the duodenum (B), the jejunum (C), and the ileum (D). Each of these compartments is made up of two glass jackets containing a flexible silicone membrane, which, through its relaxation and compression, generates the peristaltic movements. These movements are regulated by the opening and closing of the valves connecting the compartments, which allow proper mixing of the chyme. All digestion parameters (pressure, temperature, pH, and content volume) can be continuously monitored, thanks to the sensors installed in each compartment. Heated water is continuously pumped into the 4 compartments to ensure physiologically relevant temperature to that of the gastrointestinal tract. Media administered into each compartment are prepared with specific formulation ratios to match the chemical and enzymatic conditions of each compartment (gastric juice (HCl), pepsin and lipase in the stomach, pancreatin solution (sodium bicarbonate) and trypsin in the duodenum and small intestinal electrolyte solution in the other compartments). Jejunum and ileum are connected to hollow filter membranes (11.8 kDa molecular weight cut-off, Baxter Xenium 110 high-flux dialyzer, Baxter Healthcare, Deerfield, US) simulating an absorptive step. *Text adapted from Barker R. et al., J. Pharm. Sci.*, *2014*.

**Supplementary Fig. S4.**
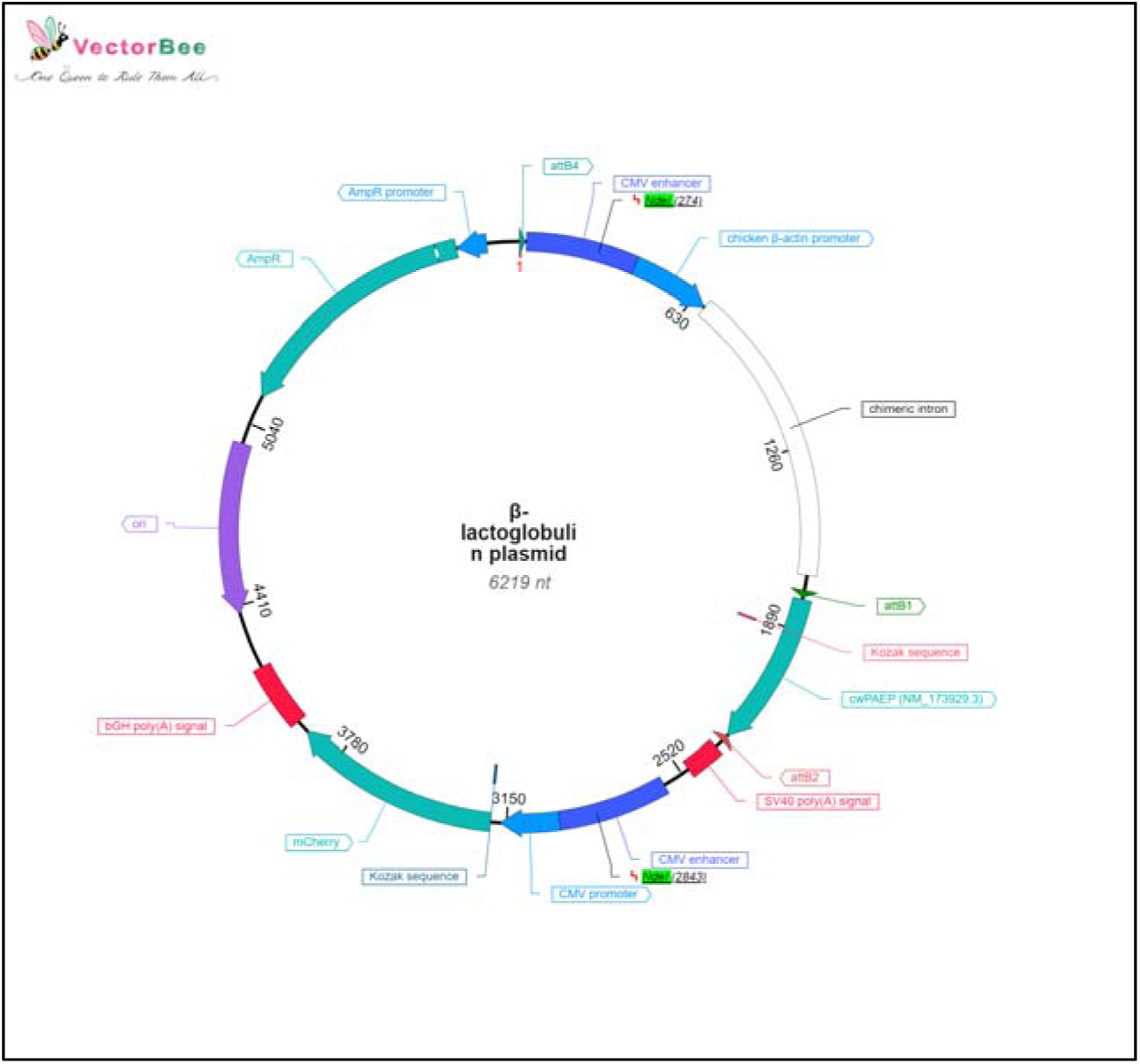
β-lactoglobulin plasmid map. The above plasmid is a mammalian expression plasmid carrying the open reading frames coding for the following three genes: mCherry (a fluorescent protein whose signal was used to assess transfection efficiency between different conditions upon plasmid transfection into HEK293T cells), ampicillin resistance gene (AmpR, the antibiotic resistance gene used for bacterial selection during plasmid cloning) and the gene coding for β-lactoglobulin (cwPAEP). NdeI restriction sites are denoted on the map (NdeI highlighted in green). The vector was custom-designed by VectorBuilder Inc. The sequence of the plasmid was obtained from the VectorBuilder Inc. database, and the map was built using the VectorBee software 2.5.0-WIN.

**Supplementary Fig. S5.**
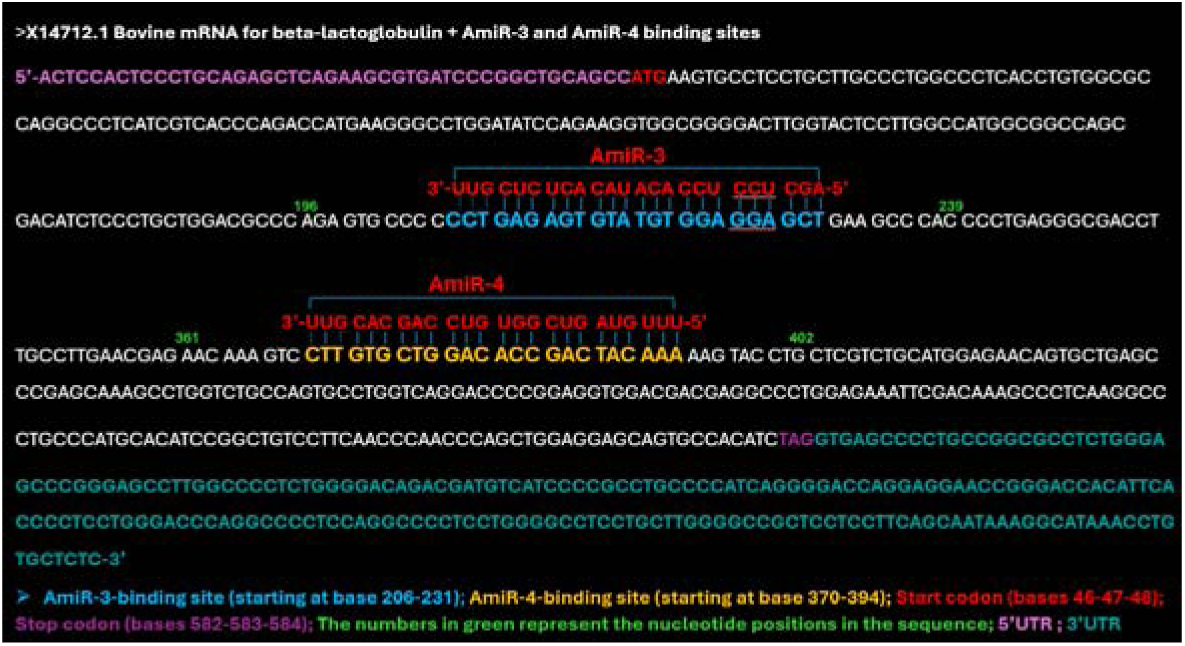
Scheme depicting the binding sites of AmiR-3 and AmiR-4 in the coding sequence of β-lactoglobulin mRNA (Genbank no. X 14712.1).

**Supplementary Fig. S6.**
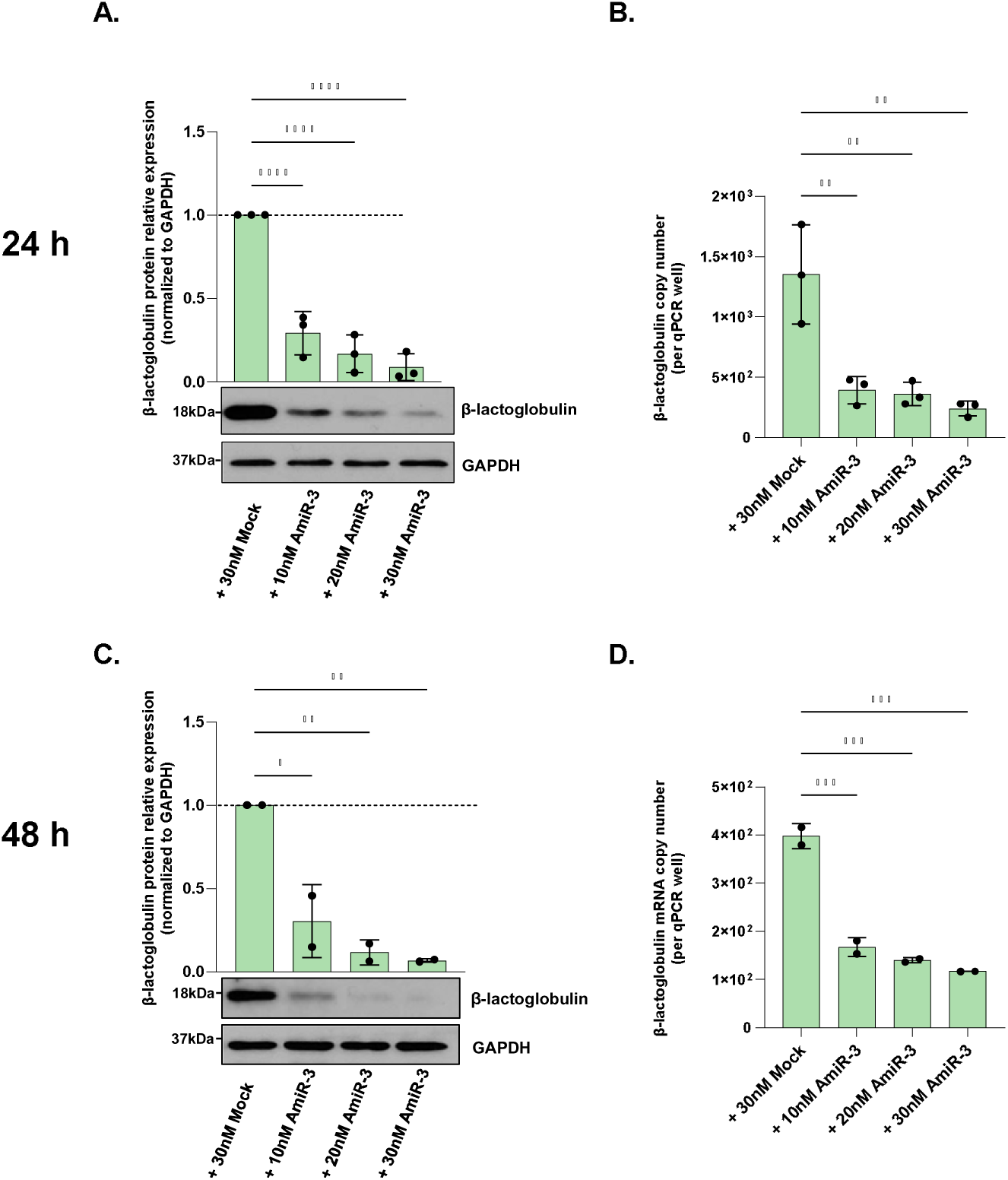
Estimation of the AmiR “effective” copy number. HEK293T cells were co-transfected with a plasmid expressing β-lactoglobulin along with different concentrations of AmiR-3/4 mimics. **A-B.** Western blots, densitometry, and RT-qPCR-based absolute quantification of β-lactoglobulin protein and mRNA levels in HEK293T cells after co-transfection with β-lactoglobulin plasmid and AmiR-3 mimic at three different concentrations (10 nM, 20 nM, and 30 nM) for 24 h (n=3). **C-D.** Western blots, densitometry, and RT-qPCR-based absolute quantification of β-lactoglobulin protein and mRNA levels in HEK293T cells after co-transfection with β-lactoglobulin plasmid and AmiR-3 mimic at three different concentrations (10 nM, 20 nM, and 30 nM) for 48 h (n=2). The microRNA mimic mock, Cel-67, was used as a negative control at 30 nM. GAPDH was used as an internal control for western blot densitometry analysis and for RT-qPCR. The values are means ± SD of three independent experiments (n=2-3). Statistical analysis was performed using one-way ANOVA followed by Tukey’s post hoc test (*p<0.05, **p<0.01, ***p<0.001, ****p<0.0001).

**Supplementary Fig. S7.**
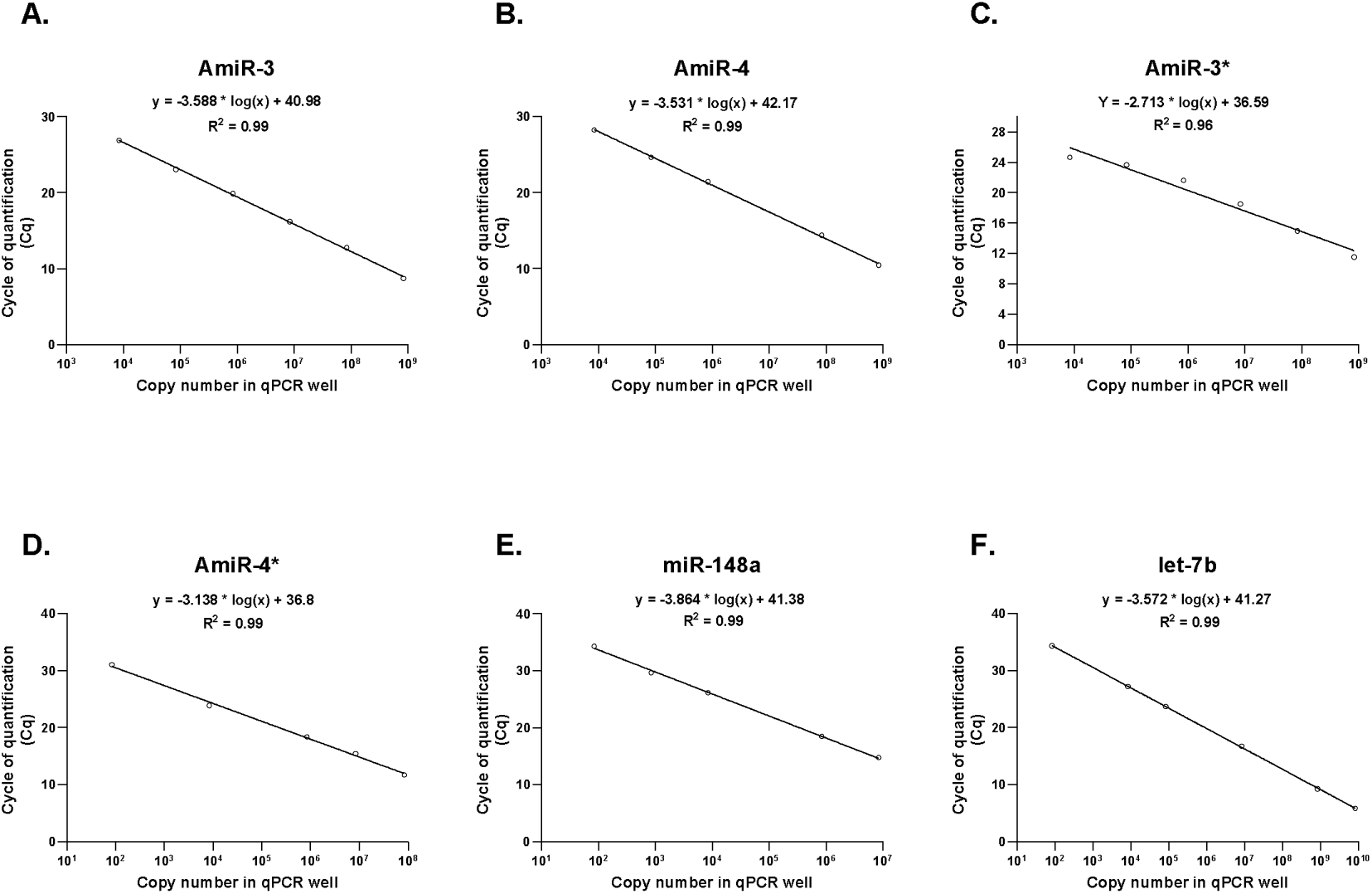
Standard curves for microRNA RT-qPCR-based absolute quantification. Standard curves showing the copy number (in RT-qPCR well) of **A**. AmiR-3, **B**. AmiR-4, **C**. AmiR-3*, **D**. AmiR-4*, **E**. miR-148a, and **F**. let-7b were established using the synthetic RNA oligonucleotides (IDT, Coralville, IA, USA) serially diluted 1/10th to obtain a concentration of 5-6 logs.

